# Adaptation of the freshwater anaerobic methanotroph ‘*Ca*. Methanoperedens vercellensis’ to low pH levels reveals membrane lipid remodelling

**DOI:** 10.64898/2026.04.11.717812

**Authors:** Vojtěch Tláskal, Reinier A. Egas, Wei Wang, Xiaoxiao Zhao, Martijn Wissink, Maider J. Echeveste Medrano, Kevin W. Becker, Felix J. Elling, Cornelia U. Welte

## Abstract

Anaerobic methanotrophic archaea are key members of the biological methane filter, thereby preventing emissions of this strong greenhouse gas into the atmosphere. Previous studies on freshwater anaerobic methanotrophs targeted the activity of these microorganisms at circumneutral pH whereas molecular ecology studies identified this phylotype also in acidic environments such as peatlands; it is currently unknown whether they can adapt to low pH and remain effective in the biological methane filter in low pH environments. Here we show that a granular enrichment culture of the freshwater methanotroph ‘*Ca*. M. vercellensis’ loses activity when experiencing pH stress but remains metabolically active down to pH 5.65 with appropriate adaptation time, indicating that adaptive changes are necessary to accommodate anaerobic methane oxidation at lower pH. Analyses of archaeal lipids revealed an increase in zwitterionic intact polar lipids over anionic lipids as an adaptation. This coincided with a change in granule structure while methane oxidation rate and enrichment state of ‘*Ca*. M. vercellensis’ remained stable. We show that ‘*Ca*. M. vercellensis’ remains metabolically active at lower pH values, despite increased maintenance energy demands and the need for cytoplasmic pH homeostasis. Our study demonstrates that adaptations to stress by slow-growing microorganisms may require long-term observation and is thereby instrumental for a better understanding of methane cycling in acidic ecosystems.

## Introduction

Methane (CH_4_) represents the second most abundant greenhouse gas, with a 27-fold stronger warming potential than CO₂ on a 100-year timescale (IPCC, 2023). Atmospheric methane levels have increased 2.6 times (to 1.91 ppmv in 2022) compared to pre-industrial value (Lan et al., 2025). While anthropogenic emissions represent a substantial portion of total methane, large quantities are emitted from wetlands and inland freshwater systems (Saunois et al., 2025). Methane is produced by methanogens in anoxic environments with very low redox potentials, such as sediments and gastrointestinal tracts. Anaerobic oxidation of methane (AOM) represents an important biological sink of methane and is performed by anaerobic methane-oxidising (ANME) archaea. These microbes are able to use reverse methanogenesis to oxidise methane using a variety of electron acceptors such as sulfate, metal oxides, organic acids, and nitrate (N-DAMO) (Boetius et al., 2000; Raghoebarsing et al., 2006; Haroon et al., 2013; Ettwig et al., 2016; Welte et al., 2016; Cai et al., 2018; Leu et al., 2020; Valenzuela and Cervantes, 2021). ANME archaea are an integral part of the biological methane filter that partly offsets methane emissions from natural habitats. Their activity has been described in marine sediments (Boetius et al., 2000; Wallenius et al., 2025), freshwater sediments (Timmers et al., 2016; Echeveste Medrano et al., 2025b), and wetlands (Valenzuela et al., 2017; Krause and Treude, 2021). However, the range of environmental conditions at which ANME archaea thrive has not yet been fully explored.

Members of the archaeal family *Methanoperedenaceae* (formerly ANME-2D, phylum Halobacteriota), with the genus ‘*Candidatus* (*Ca*.) Methanoperedens’ are key methane oxidisers in anoxic freshwater environments (Welte et al., 2016; Liu et al., 2026). Salinization from ongoing sea level rise threatens freshwater habitats, while eutrophication can lead to increased sulfide concentrations. ‘*Ca*. Methanoperedens’ has been tested for adaptation to changes in salt concentration and sulfide stress (Echeveste Medrano et al., 2024, 2025a). Accumulation of storage polymers in the cells of ‘*Ca*. Methanoperedens’ contributes to cell stability under osmotic stress (Frank et al., 2023; Echeveste Medrano et al., 2024). Currently, there are limited data available as to how anaerobic methanotrophs react to pH decrease and acid stress in freshwater, wetland, and peatland environments, where widespread AOM has been observed (Smemo and Yavitt, 2011; Gupta et al., 2013; Segarra et al., 2015; Barney et al., 2024). Peatlands with limited buffering capacity develop strongly acidic conditions (pH 3.5–4.5) as a result of organic-matter decomposition and atmospheric deposition. Such low pH can markedly affect the physiology and cellular adaptations of anaerobic methanotrophs (Gupta et al., 2013; Miller et al., 2019; Barney et al., 2024; Nweze et al., 2024). Moreover, peat-soil pH is modulated by seasonal fluctuations, drainage, afforestation-induced acidification, and acid deposition, each imposing additional selective pressures that require methanotrophs to adjust their metabolic traits (Malcolm et al., 2014; Urbanová and Bárta, 2016; Kang et al., 2018; Harpenslager et al., 2024).

Studying these responses is complicated by the experimental constraints associated with ANME research; namely that isolation of ‘*Ca*. Methanoperedens’ is challenging owing to its slow doubling times (Timmers et al., 2017) and lack of axenic cultures. Enrichment bioreactor systems enable physiological studies (McIlroy et al., 2023; Echeveste Medrano et al., 2024, 2025a; Liu et al., 2026) that have to be interpreted with care due to the presence of a side community consisting of nitrite-utilising autotrophs, which detoxify nitrite, and heterotrophs thriving off necromass and syntrophic intermediary products (Cai et al., 2019). An advanced enrichment of ‘*Ca*. Methanoperedens’ has been achieved by using electrodes in bioelectrochemical systems as electron acceptors instead of nitrate (Ouboter et al., 2022, 2024). Biofilm enrichment cultures, however, show high sensitivity to a decrease of pH.

Potential adaptation to pH stress in archaea is represented by the ability to modulate the composition of lipids in the archaeal membrane. In environments with high temperature and low pH, archaea modulate membrane fluidity and permeability by synthesizing glycerol dibiphytanyl glycerol tetraether (GDGT) lipids with cyclopentane rings (Zhou et al., 2020). A higher degree of GDGT cyclization leads to tight packing of tetraether lipids which restricts proton influx into the cytoplasm (Chong, 2010). A negative correlation of pH and ring index was indeed described in several thermoacidophilic archaeal taxa such as in *Acidilobus* (Boyd et al., 2011), *Picrophilus* (Feyhl-Buska et al., 2016), or *Sulfolobus* (Cobban et al., 2020) and in hot springs (Boyd et al., 2013). Likewise, the abundance of cyclization genes *grsAB* for GDGT ring synthase A and B (Zeng et al., 2019) was found to be negatively correlated with pH in hot spring metagenomes (Blum et al., 2023). In terms of acid-adaptation at the level of intact polar lipids (IPLs), Shimada et al. (2008) reported an increase in the number of sugar units in glycophospholipids of the thermoacidophilic archaeon *Thermoplasma acidophilum*. The only study on the influence of pH on intact polar and core lipid adaptation in archaea living at moderate temperatures observed no change in ring index, a small decrease in anionic IPLs and a corresponding increase in IPLs with two sugar units upon decreases in pH within the circumneutral range (Elling et al., 2015). However, membrane lipid adaptation in archaea living in moderate temperature and acidic pH conditions, such as in many wetlands, has not been studied. Membrane lipids of ‘*Ca*. Methanoperedens carboxydivorans’ (formerly known as BLZ2, Arshad et al., 2015; Berger et al., 2017) and ‘*Ca*. Methanoperedens vercellensis’ have been profiled showing archaeol and hydroxyarchaeol as the main core lipids and dihexose as the main component of IPL headgroups (Kurth et al., 2019). The ability of ‘*Ca*. Methanoperedens’ to adapt to acid stress is, however, unknown.

Based on the environmental diversity of the genus ‘*Ca*. Methanoperedens’, we investigated whether an enrichment culture of ‘*Ca*. M. vercellensis’ can adapt to a stepwise decrease in pH in a continuous laboratory-scale bioreactor. The use of ^13^C-CH_4_ as an energy and carbon source (Kurth et al., 2019) allowed assessment of the methane oxidation potential of the enrichment culture under acid stress. Optical microscopy and fluorescent *in situ* hybridisation (FISH) were employed to examine the morphology of ‘*Ca*. M. vercellensis’ granular biomass. Furthermore, analysis of archaeal lipids and analysis of genes encoding biosynthetic proteins involved in cell membrane composition remodelling were used to investigate the mechanism of ‘*Ca*. Methanoperedens’ adaptation to low pH.

## Methods

### Bioreactor setup and operation

A 6.5 L bioreactor (Applikon, Delft, The Netherlands) with 4.6 ± 0.0 L of liquid phase was anoxically inoculated with 100 mL biomass of a ‘*Ca.* Methanoperedens vercellensis’ enrichment originating from freshwater Italian rice paddy fields (Vaksmaa et al., 2017; Echeveste Medrano et al., 2024; Wissink et al., unpublished data). The bioreactor was operated as a sequencing fed-batch reactor (SBR) resulting in granular biomass for 172 days, after which the acidification experiment started and lasted for 246 days. The bioreactor operated on a daily cycle without a settling phase before supernatant removal to stimulate granule formation. Each cycle consisted of continuous medium supply (828 ± 93 mL day^-1^) followed by 45 min of supernatant removal, resulting in a hydraulic retention time of approximately 6.6 days. The bioreactor contained two standard six-blade turbines stirring at 200 rpm and was operated at room temperature. Stirring was stopped during supernatant removal to prevent granular biomass washout. The bioreactor was constantly fed with methane with a flow of 10 mL min^-1^ and sparged with Ar:CO_2_ (95:5). The pH was buffered with 1 M KHCO_3_ or HCl solution and controlled by a BL 931700 pH controller Black Stone (Hanna Instruments, Rhode Island, USA). The amount of NaNO_3_ substrate was on average 0.70 ± 0.13 mmol day^-1^ L^-1^ (Supplementary Figure 1). The experiment was initiated at the standard operating pH of 7.25 (Echeveste Medrano et al., 2024) and was terminated at pH 5.65 with the acidification step of 0.1 set approximately every 7 days (Supplementary Figure 2).

### Bioreactor batch activity assays

To determine the methane oxidation rate of the bioreactor microbial community, the bioreactor was run as a batch with 10% ^13^C-CH_4_ and additional Ar:CO_2_ (95/5) in the headspace to obtain an overpressure of 0.18–0.2 bar. The headspace was first flushed for > 1 h with Ar:CO_2_ (95/5) at 40 mL min^-1^ to remove residual methane traces. The starting nitrate concentration was ≈2 mM. Over a period of 5–7 days, labelled ^13^C-CO_2_ and ^12^C-CO_2_ were measured, and nitrate limitation, overpressure, and a stable pH were controlled. Gas concentrations were measured using 50 μL of bioreactor headspace samples (n = 2) at each timepoint by gas chromatography–mass spectrometry (GC-MS) using an Agilent 8890 GC System (Agilent Technologies, USA). Calibration was performed with standard gas consisting of He/O_2_/N_2_/CH_4_/CO_2_/N_2_O with terminal values of (%): balance/1.02/1.03/1.05/1.04/0.050 (Linde Gas Benelux BV, The Netherlands). Ratio of labelled ^13^C-CO_2_ and ^12^C-CO_2_ was used as a proxy for ongoing anaerobic oxidation of methane. Methane oxidation rate was assessed by fitting linear regression to the ^13^C-CO_2_:^12^C-CO_2_ increase as a function of time (R^2^ = 0.88 ± 0.04). Activity assays were normalized by dry weight (n = 2, 15 mL). At the end of each batch activity assay, 15 mL of bioreactor biomass were sampled, spun at 3,000 rpm for 5 min, supernatant was removed and pelleted biomass was immediately stored at -20°C.

### Nucleic acid extractions and ddPCR

DNA was extracted from the frozen bioreactor biomass pellets using a phenol-chloroform extraction method (Angel et al., 2021). For each extraction, 2 mL of bioreactor enrichment culture equivalent were used. The proportion of ‘*Ca*. Methanoperedens’ cells to the rest of the prokaryotic community members in the enrichment culture was determined using droplet digital PCR (ddPCR) on a Bio-Rad QX200 Droplet Digital PCR System. Quantification of the ‘*Ca*. Methanoperedens’ cells was assessed by targeting the *mcrA* gene with primers mcrA_rev (5’-CGTTCATBGCGTAGTTVGGRTAGT-3’, (Steinberg and Regan, 2008)) and mlas_mod (5’-GGYGGTGTMGGDTTCACMCARTA-3’, (Angel et al., 2011)). The proportion of the side community members was assessed using general 16S rRNA gene primers BAC338F (5’-ACTCCTACGGGAGGCAG-3’) and BAC805R (5’-GACTACCAGGGTATCTAATCC-3’, (Yu et al., 2005)). ddPCR premix for *mcrA* or 16S rRNA gene quantification contained 11 μL of QX200™ ddPCR™ EvaGreen Supermix, 0.2 μL of each primer (0.1 μM), 8.6 μL of H_2_O and 2 μL of template DNA. PCR conditions of gene amplification were 5 min at 95 °C, 5 cycles of 2.5 min at 60 °C with 1 °C decrease at each cycle, 34 cycles of (30 s at 95 °C, 30 s at 55 °C and 2 min at 60 °C) and 5 min at 4 °C followed by 5 min at 90 °C (Tian et al., 2026). The non-specificity of the primers for the 16S rRNA gene was confirmed by in-silico annealing to the closed genome of ‘*Ca.* Methanoperedens carboxydivorans’ (Egas et al., 2025). The genome contains a single copy of the *mcrA* gene (accession PRJEB98224).

### AOM inhibition in an activity assay

To link methane oxidation to ‘*Ca.* Methanoperedens vercellensis’, a batch activity assay with the AOM inhibitor 3-bromopropanesulfonic acid (3-BPS) was performed. 3-BPS is a structural analogue to the 2-bromoethanesulfonic acid sodium salt (2-BES). Both are MCR inhibitors and were shown to inhibit the activity of anaerobic oxidation of methane by ‘*Ca.* M. carboxydivorans’ (Wissink et al., 2024). 40 mL aliquots of granular biomass were withdrawn anoxically from the bioreactor (pH = 6.36, experiment day 148) and washed three times in medium through settling followed by liquid removal. The biomass was resuspended in triplicated 120 mL serum flasks in a final medium volume of 40 mL. After closing the serum flasks with red butyl rubber stoppers (Terumo, Leuven, Belgium), the headspace was flushed with N_2_ for 5 min, after which the pressure was set to 1.5–1.6 bar. Medium was supplemented with nitrate to the initial concentration of 1 mM and 3-BPS was added to half of the bottles to reach a final concentration of 20 mM (Wissink et al., 2024). CO_2_ (0.7% vol/vol final) and ^13^C-CH_4_ (99% stock, 3.3% final vol/vol) were added to the headspace of each flask. pH was adjusted using anoxic HCl and KOH to the final value of 6.49 ± 0.1. Positive controls consisted of medium containing biomass, methane, and nitrate. Flasks were incubated at 30 °C with shaking at 200 rpm. Following a 1-hour pre-incubation period, the headspace was analysed regularly over 14 days by GC-MS as described above. The pH was monitored and adjusted as needed throughout the incubation period. Nitrate and nitrite concentrations were measured at regular intervals. To prevent toxic nitrite accumulation, nitrate was replenished to a final concentration of 1 mM based on the consumption rate.

### Short-term response batch activity assays

The short-term tolerance of ‘*Ca.* M. vercellensis’ to acidic conditions was tested in batch bottle activity assays. As inoculum, 40 mL aliquots of granular biomass were anoxically withdrawn from ‘*Ca.* M. vercellensis’ bioreactor (Ouboter et al., 2024). Aliquots were transferred to an anaerobic chamber, washed and resuspended in 40 mL of medium in 120 mL serum flasks as described above. Medium was supplemented with nitrate to an initial concentration of 2 mM. A bacteria-suppressing antibiotic cocktail containing streptomycin, vancomycin, ampicillin, and kanamycin was added to reach a final concentration of 50 μg mL^-1^ each (Wissink et al., 2024). CO_2_ (2% vol/vol final) and ^13^C-CH_4_ (99% stock, 10% final vol/vol) were added to the Ar-filled headspace of each flask with pressure adjusted to 1.2 bar. To buffer pH, 4-(2-hydroxyethyl)-1-piperazineethanesulfonic acid (HEPES) buffer at pH 7.25 was used, pH was adjusted using anoxic HCl and KOH to the final value of pH 7.25, pH 6.25, and pH 5.65 (n = 3 per pH level). Flasks were incubated at 30 °C with shaking at 200 rpm. Following a 2.5-hours pre-incubation period, the headspace was analysed regularly over 95 hours by GC-MS as described above with replicated GC-MS injections (n = 2 per bottle per timepoint). Nitrate and nitrite concentrations were measured at regular intervals and nitrate was replenished based on the consumption rate. Methane oxidation rate was assessed by fitting linear regression as described above (R^2^ = 0.85–0.88). Activity assays were normalized by dry weight obtained from 40 mL bioreactor aliquots (n = 3).

### Intact polar lipid analysis

Lipids were extracted using a modified Bligh & Dyer protocol (Bligh and Dyer, 1959; Sturt et al., 2004), dried under a flow of N_2_, and reconstituted in methanol for analysis. Lipids were analyzed in a two-step procedure. First, lipids were identified by ultra-high performance liquid chromatography (UHPLC) coupled to high-resolution mass spectrometry (MS). These identifications were used as a reference for lipid identification and quantification of all other samples using a low resolution HPLC-MS system. The UHPLC-MS system consisted of a Vanquish Horizon UHPLC equipped with a Waters Acquity BEH C_18_ column (2.1 × 150 mm, 1.7 μm particle size) coupled to a Q Exactive Plus high-resolution, accurate mass Orbitrap MS (Thermo Fisher Scientific, Bremen, Germany) equipped with a heated electrospray ionization source (HESI-II) operating in positive ionization mode. UHPLC-MS reversed-phase chromatographic conditions were as described previously (Wörmer et al., 2013). In brief, the injection volume was 10 µL, and lipids were eluted at a constant flow rate of 0.4 mL min^-1^ using linear gradients of methanol-water (85:15, v/v; eluent A) to methanol-isopropanol (50:50, v/v; eluent B), both with 0.04% formic acid and 0.1% NH_3_. The initial condition was 100% A held for 2 min, followed by a gradient to 15% B in 0.1 min and a gradient to 85% B in 18 min. The column was then washed with 100% B for 8 min. The column temperature was 65 °C. HESI and MS settings were optimized by infusion of a mixture of lipids into the eluent flow from the LC system using a T-piece. HESI-II settings were: capillary temperature, 250 °C; sheath gas (N_2_) pressure, 30 arbitrary units (AU); auxiliary gas (N_2_) pressure, 10 AU; spray voltage, 3.5 kV; probe heater temperature, 350 °C; S-lens 75 V. Lipids were detected by scanning from *m/z* 150 to 2000, followed by data-dependent MS^2^ (isolation window 1.2 *m*/*z*) using resolving powers of 140,000 and 17,500 (FWHM at *m/z* 200) in full-scan MS and MS^2^, respectively. Optimal fragmentation of lipids was achieved using a stepped normalized collision energy of 15, 25, and 50 eV. A mass calibration was performed using the Thermo Scientific Pierce LTQ Velos ESI Positive Ion Calibration Solution prior to analysis. Lipids were identified by retention time, accurate molecular mass, and MS^2^ fragment spectra as described previously (Yoshinaga et al., 2011).

The HPLC-MS system consisted of an Agilent Series 1260 Infinity high-performance liquid chromatograph (HPLC) coupled to an Agilent 6130 single quadrupole mass spectrometer (MS). The MS was used in scan mode (200–2000 *m*/*z*) and was equipped with an ESI source operating in positive mode. The HPLC was equipped with an ACE3 C_18_ column (2.1 × 150 mm, 3 µm particle size, Advanced Chromatography Technologies, Aberdeen, Scotland). Chromatographic conditions were as described previously (Zhu et al., 2013). In brief, lipids were eluted at a flow rate of 0.2 mL min^-1^ at 45 °C. The HPLC program was as follows: 100% eluent A (methanol:formic acid:14.8 M NH_4_OH, 100:0.04:0.10, v/v/v) for 10 min, followed by a linear gradient to 24% eluent B (2-propanol:formic acid:14.8 M NH_4_OH, 100:0.04:0.10, v/v/v) in 5 min, followed by a gradient to 65% B in 55 min. The column was then flushed with 90% B for 10 min and re-equilibrated with 100% A for 10 min. ESI settings were optimized using extracts of *Methanosarcina mazei* Gö1 and *Sulfolobus acidocaldarius* DSM639 and were as follows: drying gas (N_2_) temperature at 350 °C, drying gas flow rate at 6 L min^-1^, nebulizer pressure at 35 psi, capillary voltage at 3500 V. Lipids were identified by molecular mass and retention time. Due to a lack of authentic standards, lipid abundances should be considered as semi-quantitative.

### Core lipid analysis

Core lipid extracts were prepared by direct acid hydrolysis of biomass (Elling et al., 2014). In brief, cell pellets were dissolved in 3 M HCl in methanol by vortexing (30 s) and ultrasonication (5 min) and then heated for 3 h at 70 °C. After cooling to room temperature, ultrapure water was added and the lipids were extracted three times using equal amounts of dichloromethane. Core lipids were analyzed using the HPLC-MS system described above but equipped with an atmospheric pressure chemical ionization interface operating in positive mode. Source conditions were optimized using extracts of *Sulfolobus acidocaldarius*. The source conditions were as follows: drying gas (N_2_) temperature at 200 °C, vaporizer temperature at 400 °C, drying gas flow rate at 6 L min^-1^, nebulizer pressure at 60 psi, capillary voltage at 3500 V, and corona current at 5.0 µA. Lipids were separated at a flow rate of 0.2 mL min^-1^ using two Waters Acquity BEH HILIC columns (2.1 × 150 mm, 1.7 µm particle size in series maintained at 30 °C, preceded by a guard column (2.1 × 5 mm) of the same material, as described by (Hopmans et al., 2016). The following linear gradient was used with eluents A (*n*-hexane) and eluent B (*n*-hexane:isopropanol, 90:10, v/v): 82% A for 25 min (isocratic), to 35% B in 25 min and finally to 100% B in 30 min. In variation to the method described by Hopmans et al. (2016), the column was flushed using 100% B for 10 min and equilibrated to starting conditions with 18% B for 20 min (Roy et al., 2025).

To detect GDGT ring synthase genes *grsAB* in the genome of ‘*Ca.* M. vercellensis’, amino acid sequences of enzymes GrsA and GrsB from a model organism *Sulfolobus acidocaldarius* (Zeng et al., 2019) were retrieved from Uniprot (Q4J8I0 and Q4JC22 for GrsA and GrsB, respectively). ‘*Ca.* M. vercellensis’ genes were predicted with Prokka v1.14.6 (Seemann, 2014) and blastp v2.15.0+ was used to search for candidate *grsAB* genes.

### Fluorescence *in situ* hybridization and optical microscopy

Biomass (15 mL) was sampled from the bioreactor before the start of the acidification experiment (39 days before, pH = 7.25) and at the end of the experiment (after 246 days, pH = 5.65). Biomass samples were immediately pelleted, supernatant was removed, the cell pellet was resuspended in 500 μL PBS and mixed with 1500 μL of 4% paraformaldehyde. The sample was incubated for 3 h at 4 °C and subsequently washed three times with PBS. Washed biomass was resuspended in 1 mL of 1:1 PBS:ethanol (97%) and stored at -20 °C until FISH analysis. FISH was performed as described by Wagner et al. (2003) on individual granules from the bioreactor. PFA-fixed granules were transferred onto a 10-wells slide and dried for 15 min at 46 °C. The slide was dehydrated by dipping in a graded ethanol series for 3 min each (50%, 80%, 100%) and dried at 46 °C. Then 10 µL of hybridization buffer was added to the sample followed by 1 µL of each probe. The hybridization buffer contained (per 1 mL): 400 µL formamide, 399 µL Milli-Q water, 20 µL 1 M Tris-HCl, 180 µL 5 M NaCl and 1 µL 10% SDS. The probe mix contained: Arch915, 5′-GTGCTCCCCCGCCAATTCCT-3′ (Stahl, 1991) (Fluos; 30 ng µL^−1^) targeting archaea; EUB338-I, 5′-GCTGCCTCCCGTAGGAGT-3′ (Amann et al., 1990) (Cy5; equimolar at 50 ng µL^−1^) targeting bacteria. A nonsense probe NON-EUB not targeting prokaryotic rRNA (5′-ACTCCTACGGGAGGCAGC-3′, Wallner et al., 1993) was applied to samples as a negative control for binding, hybridization carried out without any probe served as a control for autofluorescence. Probe hybridization occurred at 46 °C in a humid chamber for 20 h. The slide was then immersed in a wash buffer at 48 °C for 15 min, followed by brief immersion in ice-cold Milli-Q water and air drying. The wash buffer contained (per 50 ml): 0.46 mL 5 M NaCl, 1 ml 1 M Tris-HCl, 0.5 mL 0.5 M EDTA, and 48 mL Milli-Q water. Dried slides were stored at -20 °C until microscopy analysis. Biomass granules were mounted and counterstained with 10 µL of ProLong Diamond Antifade Mountant with DAPI (Thermo Fisher Scientific). Cells were analysed by confocal microscopy with a FV3000 Laser Scanning Microscope (Olympus, Japan) at the Light Microscopy Core Facility, BC CAS.

For optical microscopy of granules, pictures of thawed biomass from bioreactor with the enrichment culture of ‘*Ca.* M. vercellensis’ were taken on an optical microscope with a millimeter scale bar. Granules of the enrichment culture grown at pH 7.25 and pH 5.65 were analysed.

### Statistics

All statistical analyses and visualization were performed using the R statistical language versions 4.3.3 and (R Core Team, 2024) and with the packages tidyverse v2.0.0 (Wickham et al., 2019), tidylog v1.1.0 (Elbers, 2020), broom v1.0.9 (Robinson and Hayes, 2023), readr v2.1.5 (Wickham et al., 2024), readxl v1.4.5 (Wickham and Bryan, 2023), model fitting was performed using base stats and model parameters were displayed using report package v0.6.1 (Makowski et al., 2023). Mean values ± standard errors are given for the groups. Values of p ≤ 0.05 were considered statistically significant.

## Results and Discussion

### Long-term pH stress in a laboratory-scale bioreactor yields a stable and active methane oxidising culture while short-term stress inhibits methane oxidation

To assess whether ‘*Ca*. M. vercellensis’ can tolerate short-term acid stress without prior adaptation, 5-day batch activity assays were conducted at three pH levels. While the biomass at pH 7.25 was actively oxidizing methane (0.062 ± 0.009 change in ^45^CO_2_:^44^CO_2_ g_DM_ day^-1^, n = 3), methane oxidation rate at pH 6.25 was almost entirely inhibited (0.003 ± 0.0 change in ^45^CO_2_:^44^CO_2_ g_DM_ day^-1^, n = 3). Biomass at pH 5.65 did not show any methane oxidation activity (Figure 1).

**Figure 1.**
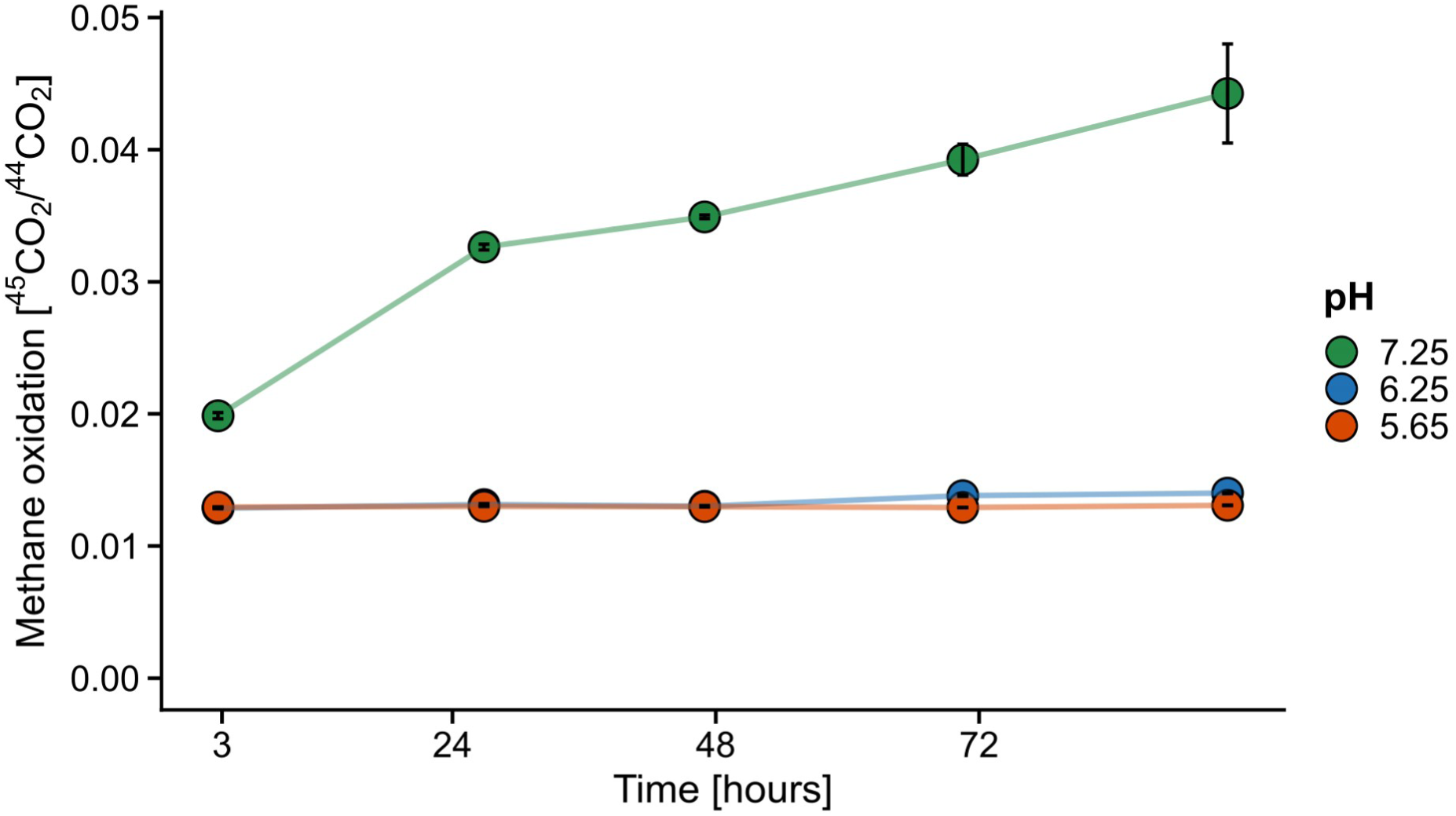
Short-term response ‘*Ca*. Methanoperedens vercellensis’ to acid stress. ^45^CO_2_/^44^CO_2_ ratio in the headspace of batch activity assay incubated at pH 7.25, pH 6.25, and pH 5.65 was used to assess the ability to oxidise ^13^C-CH₄. The headspace composition of activity assays was analysed during 95 hours of incubation. Points denote average values ± SE (n = 3 biological replicates).

We then assessed whether ‘*Ca*. M. vercellensis’ could adapt to acid stress during a gradual lowering of the pH during cultivation in a bioreactor (decrease of ∼0.1 pH units per week) with continuous medium and substrate supply. During the 225 days of stepwise acidification of the bioreactor medium, the biomass content decreased from 2.33 g L^-1^ to 0.99 g L^-1^ (57.7% decrease, Figure 2A) indicating that a part of the microbial community could not adapt to the changed pH conditions. Previous metagenome-based analyses of the parent bioreactor had indicated that ‘*Ca*. M. vercellensis’ was enriched to 54%–75% (Echeveste Medrano et al., 2024; Ouboter et al., 2024) while at the time of this study the enrichment might have been as low as 10% (Wissink et al, unpublished data). To investigate whether ‘*Ca*. M. vercellensis’ or the accompanying side community were inhibited due to pH stress, we performed quantification of the reverse methanogenesis gene *mcrA* for assessing the abundance of ‘*Ca*. M. vercellensis’ and bacterial 16S rRNA genes for quantifying the side community.

**Figure 2.**
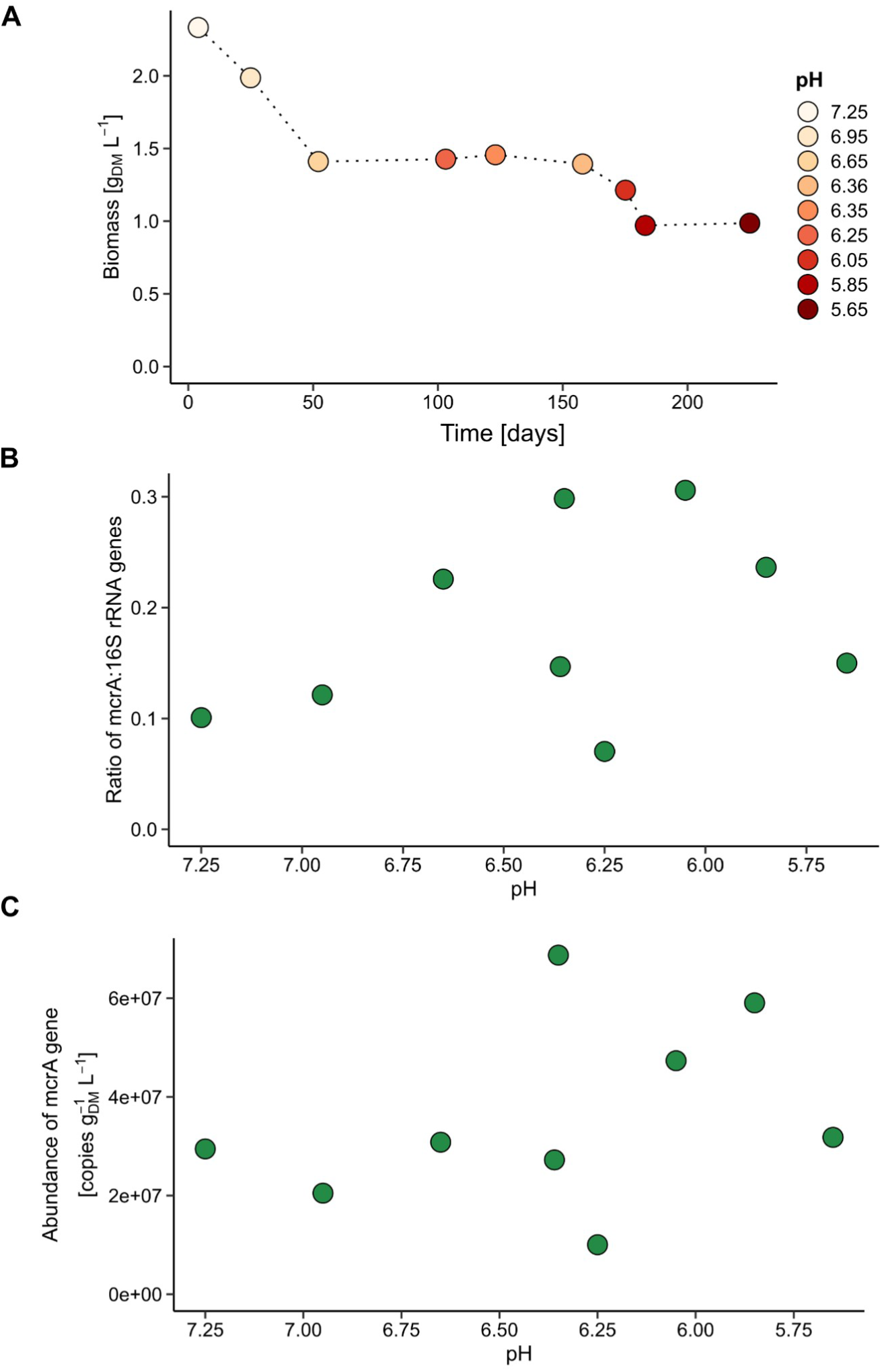
The characteristics of ‘*Ca*. M. vercellensis’ biomass in the bioreactor during the 225 days of the acidification experiment. **(A)** Decrease of dry weight of total bioreactor biomass per L of bioreactor medium. The colour of points represents pH of the bioreactor medium. **(B)** Ratio of counts of *mcrA* gene and 16S rRNA gene in DNA samples extracted from the biomass sampled at the end of individual bioreactor batch incubations. **(C)** Absolute abundance of *mcrA* gene in copies per g_DM_ per L of bioreactor medium as measured in extracted DNA.

The ratio of *mcrA* to the 16S rRNA gene showed mean values of 0.18 ± 0.03 (n = 9 , Figure 2B), indicating an enrichment of ‘*Ca*. M. vercellensis’ in the microbial community between 10%–30%, while the absolute abundance of the *mcrA* gene showed mean values of 3.61 × 10⁷ ± 6.25 × 10⁶ copies g_DM_ ^-1^ L^-1^ (n = 9, Figure 2C). Both *mcrA*:16S rRNA gene ratio and absolute *mcrA* abundance showed a non-significant negative correlation with pH (p > 0.05). The stable *mcrA* proportion and its absolute abundance together with the observed decline in total bioreactor biomass content point towards a persistence of ‘*Ca*. M. vercellensis’ in the bioreactor, with a possible enrichment towards the end of the bioreactor experiment. Furthermore, the methane oxidation rate (expressed by dry mass corrected change of ^45^CO_2_:^44^CO_2_ ratio in the headspace) was stable throughout the entire experiment and established that slow adaptation to lower pH enabled ‘*Ca*. M. vercellensis’ to adapt and remain active until pH 5.65, when the experiment was terminated. The values of methane oxidation rate peaked at pH 5.85 (0.0039 change in ^45^CO_2_:^44^CO_2_ g_DM_ day^-1^) and pH 6.65 (0.0021 change in ^45^CO_2_:^44^CO_2_ g_DM_ day^-1^) while their mean throughout the experiment was 0.0013 ± 0.0004 change in ^45^CO_2_:^44^CO_2_ g_DM_ day^-1^ (n = 9, Figure 3A). On day 100, the pH was decreased to 6.25 and associated methane oxidation rate measurements revealed that the methane oxidation activity had dropped compared to earlier measurements, so the pH was increased to pH 6.35 to allow for a prolonged adaptation period before the pH was lowered again. After ∼60 days of recovery at pH 6.35 the bioreactor had recovered its methane oxidation activity which permitted further lowering of the pH. The oxidation activity across all samples was in direct relationship to ‘*Ca*. M. vercellensis’ cell numbers as linear regression revealed a significant positive correlation between *mcrA* gene abundance and methane oxidation rate (p < 0.05, R² = 0.29, Figure 3B), indicating that ‘*Ca*. M. vercellensis’ was responsible for the observed methane oxidation activity. Cells of ‘*Ca*. M. vercellensis’ contain a single copy of the *mcrA* gene and therefore a proportional link between total cell number and rate of oxidation is expected. To exclude the possibility of interference by other members of the bioreactor microbial community, we performed an inhibition activity assay. Microcosms with the MCR inhibitor 3-BPS (Wissink et al., 2024) showed inhibition of methane oxidation during the batch activity assay while control microcosms without the addition of 3-BPS remained active, confirming that ‘*Ca*. M. vercellensis’ was responsible for methane oxidation, and that the contribution of bacterial, non-Mcr methane oxidation was negligible in our experiments (Supplementary Figure 3). During the long-term experiment, cells of ‘*Ca*. M. vercellensis’ were under increasing proton stress as the concentration of H^+^ increased 39.8× between the starting point pH 7.25 and the end point pH 5.65. In addition to proton pressure, influx of dissolved CO_2_ into the cytoplasm at low pH represents another factor which affects already stressed cells. While at pH 7.25, 89% of CO_2_ is present as bicarbonate (HCO_3_ ^-^) and uncharged CO_2_ is represented by 11.2%, at pH 5.65 the concentration of CO_2_ increases 7.4× (83.4%). Passive diffusion of this pool of uncharged CO_2_ across cell membranes and subsequent rehydration to HCO₃^-^ + H⁺ releases protons directly to the cytoplasm, dissipating the proton motive force. As the cytoplasm has a limited buffering capacity, CO_2_ influx represents an additional cellular energy demand. Cells already facing large H⁺ pressure from the extracellular environment need additional proton pumps and chaperons to maintain cytoplasmic pH homeostasis. Despite that, our data collectively indicate that ‘*Ca*. M. vercellensis’ can adapt to long-term acidification but is reversibly inactivated by immediate acid stress.

**Figure 3.**
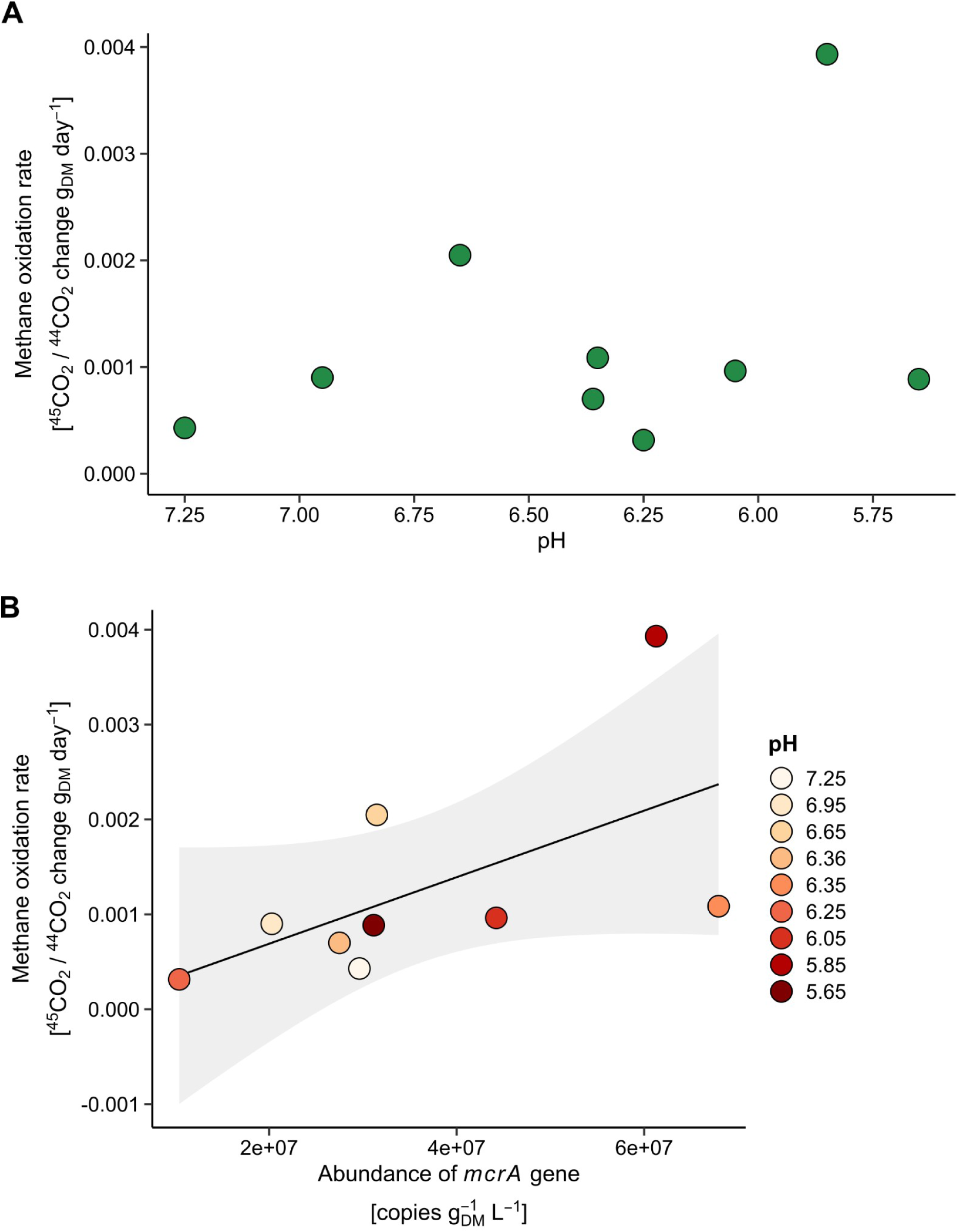
The rate of anaerobic oxidation of methane by the enrichment culture of ‘ *Ca*. M. vercellensis’. **(A)** The rate of anaerobic oxidation of methane is expressed as the slope of change in ratio of ^45^CO_2_/^44^CO_2_ in the bioreactor headspace per day and it is corrected for dry mass of the bioreactor. The measured values of the rate of anaerobic oxidation of methane during the individual bioreactor batch incubations are plotted as a function of pH. **(B)** Correlation of anaerobic oxidation of methane with the abundance of *mcrA* gene in the bioreactor biomass sampled at individual timepoints. The colour of points represents pH of bioreactor medium, solid lines indicate significant relationships (p < 0.05) with 0.95 confidence interval in grey.

### Granular biomass of ‘*Ca*. M. vercellensis’ changes in size and appearance upon adaptation to low pH

Optical microscopy of ‘*Ca*. M. vercellensis’ granules grown at pH 7.25 and pH 5.65 revealed distinct morphological differences. Granules from pH 7.25 appeared dark with black and red coloration. The red colour might be attributed to the presence of ferritin-like proteins which were observed in ‘*Ca*. M. carboxydivorans’ (Wissink et al., 2026) and the presence of *c*-type cytochromes (Arshad et al., 2015; Kletzin et al., 2015). In contrast, granules from pH 5.65 displayed white material on their surface (Figure 4). We hypothesize that this white material consists of extracellular polymeric substances (EPS) with a composition differing from that observed at pH 7.25. Additionally, acidification was associated with a reduction in granule size, consistent with the overall decline in bioreactor biomass observed over the course of the experiment (Figure 2A, 4). Similar to a previous study, fluorescence *in situ* hybridization (FISH) visualization revealed that granules consisted of ‘*Ca*. M. vercellensis’ cells accompanied by a bacterial community (Echeveste Medrano et al., 2024). Within the granules, ‘*Ca*. M. vercellensis’ formed densely packed clusters surrounded by bacterial biomass. In granules grown at pH 5.65, the ‘*Ca*. M. vercellensis’ biomass was not fully encapsulated, as portions of some clusters extended to the granule surface, becoming exposed at the outer layer (Figure 4).

**Figure 4.**
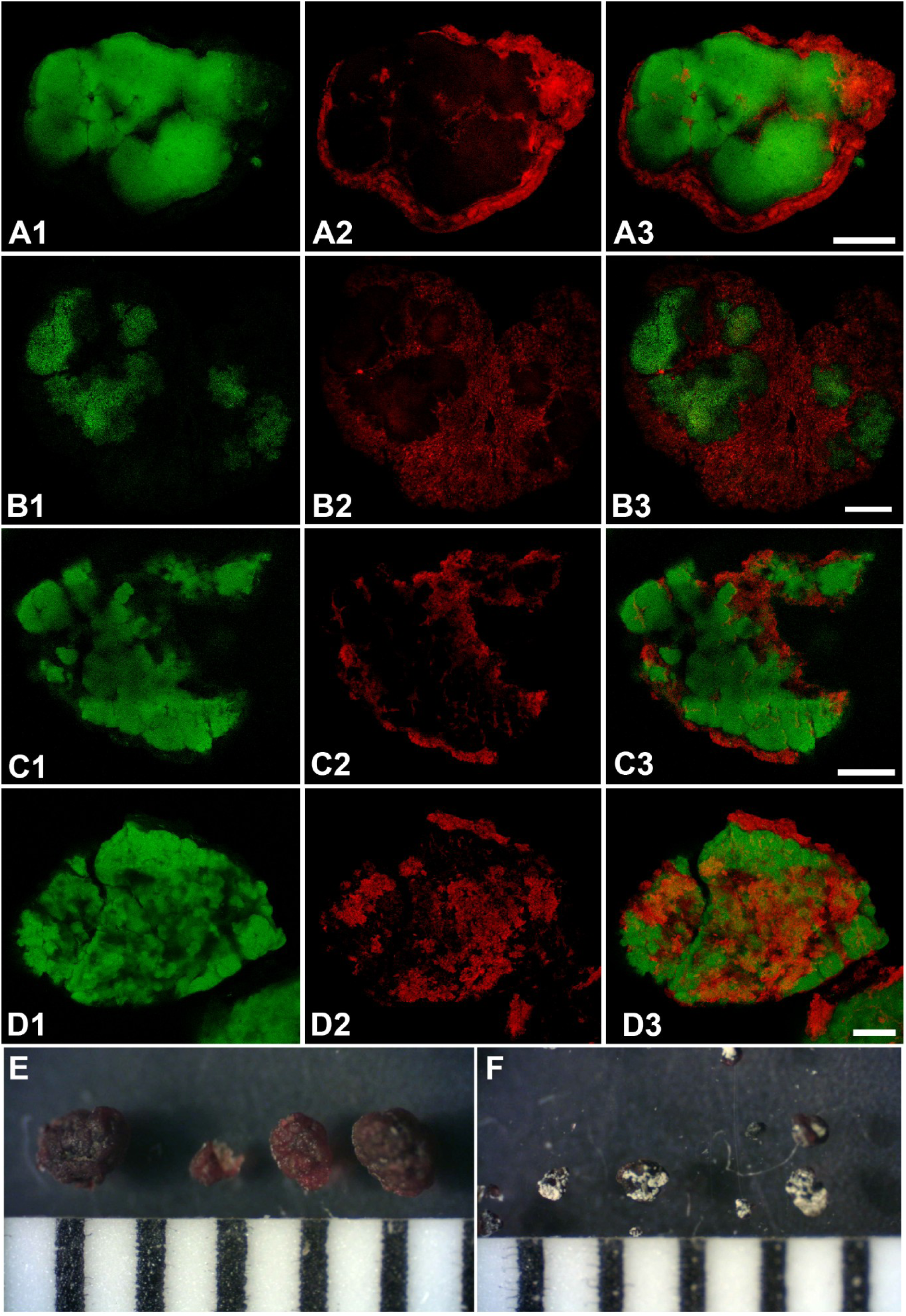
Structure, size, and composition of microbial granules from the enrichment culture of ‘*Ca*. M. vercellensis’. Composition of granules is shown in the fluorescent micrograph of biomass sample from the enrichment culture ‘*Ca*. M. vercellensis’ grown at pH 7.25 (**A, B** – one layer of a granule) and pH 5.65 (**C** – one layer of a granule**, D** – composite layers). In all fluorescent micrographs, green fluorescence labels ‘*Ca*. M. vercellensis’ (FLUOS-Arc915), whereas red fluorescence labels total bacteria (Cy5-EUBMIX) and the scale bar corresponds to 50 μm. Optical microscope picture of granules of ‘*Ca*. M. vercellensis’ grown at pH 7.25 (**E**) and pH 5.65 (**F**). The scale bar in the optical microscope picture has 1 mm steps.

### *‘Ca*. M. vercellensis’ remodels its lipid composition during adaptation to low pH

To evaluate the potential of ‘*Ca*. M. vercellensis’ to adapt to changes in pH, we analysed its genomic capacity to modify its membrane lipids as well as the chemical identity of membrane lipids of biomass originating from the bioreactor. As ‘*Ca*. M. vercellensis’ was the only archaeon in our bioreactor microbial community (Echeveste Medrano et al., 2024; Ouboter et al., 2024; Wissink et al., unpublished data) we assigned all archaeal membrane lipids to ‘*Ca*. M. vercellensis’. Many archaeal taxa have been found to protect cells from increased proton concentration in the extracellular space by producing monolayer-forming lipids (GDGTs) and by introducing cyclopentane rings into GDGTs (e.g., Boyd et al., 2011; Cobban et al., 2020). However, blastp search of the genes encoding GDGT ring synthases (*grsAB*; Zeng et al., 2019) revealed its absence from the genome of ‘*Ca*. M. vercellensis’; this was functionally confirmed by lipidomics analysis, which did not detect any GDGTs with cyclopentane rings (e.g. GDGT-1, GDGT-2) of ‘*Ca*. M. vercellensis’. Overall, GDGTs were of low abundance in this dataset. The major membrane lipids of ‘*Ca*. M. vercellensis’ were archaeols and hydroxyarchaeols, which are bilayer forming lipids. In further lipidomics analyses, we detected six core lipids (example structures are given in Figure 5A, B, C) with the most abundant being archaeol (97.27 ± 0.21%), followed by 2OH-GDGT-0 (1.00 ± 0.10%), and OH-GDGT-0 (0.62 ± 0.08%, Figure 6). Other core lipids showed a mean proportion of total lipid extract <0.5%. These three most abundant lipids also showed significant relationships with pH (Supplementary Figure 4). The archaeol proportion was positively correlated with pH (β = 0.95, p < 0.05), while 2OH-GDGT-0 and OH-GDGT-0 were negatively correlated (β = -0.44 and -0.36, respectively, p < 0.05). The observed patterns suggest a decline in archaeol content (by 1.83% overall) accompanied by an increase in hydroxylated core lipids (0.97% and 0.66% for 2OH-GDGT-0 and OH-GDGT-0, respectively) under lower pH. However, this response was quantitatively limited due to the low relative abundance of these lipids and their importance as adaptations to acid conditions may be small.

**Figure 5.**
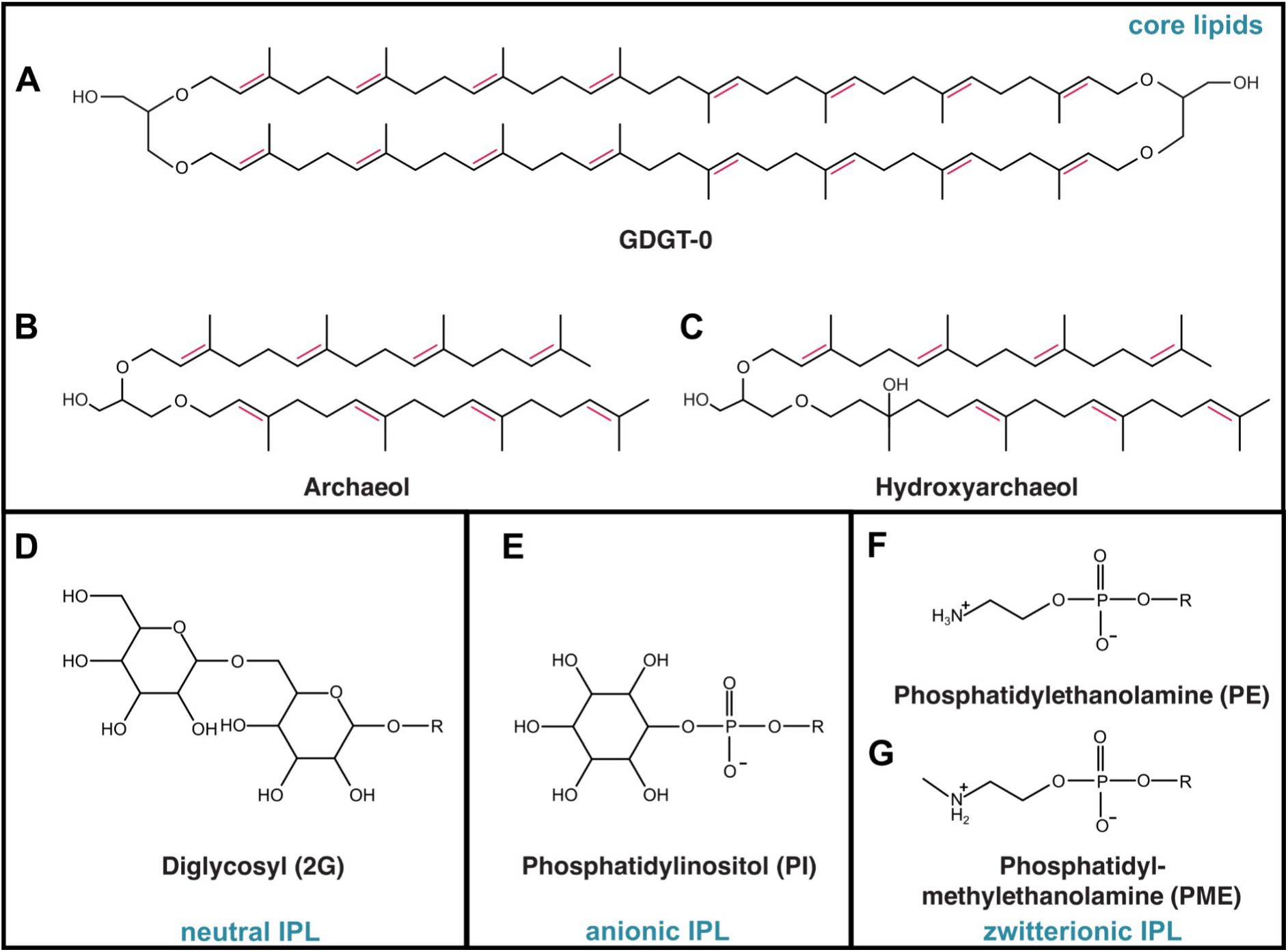
Structures of lipids identified in the biomass of ‘*Ca*. M. vercellensis’. The molecular structures of core lipids (**A, B, C**), neutral IPL (**D**), anionic IPL (**E**), and zwitterionic IPL (**F, G**) are given. Tentative positions of double bonds in core lipids are given, although the exact positions could not be resolved. Note that not all combinations of IPL headgroup, core lipid and number of core lipid unsaturations occur in ‘*Ca.* M. vercellensis’.

**Figure 6.**
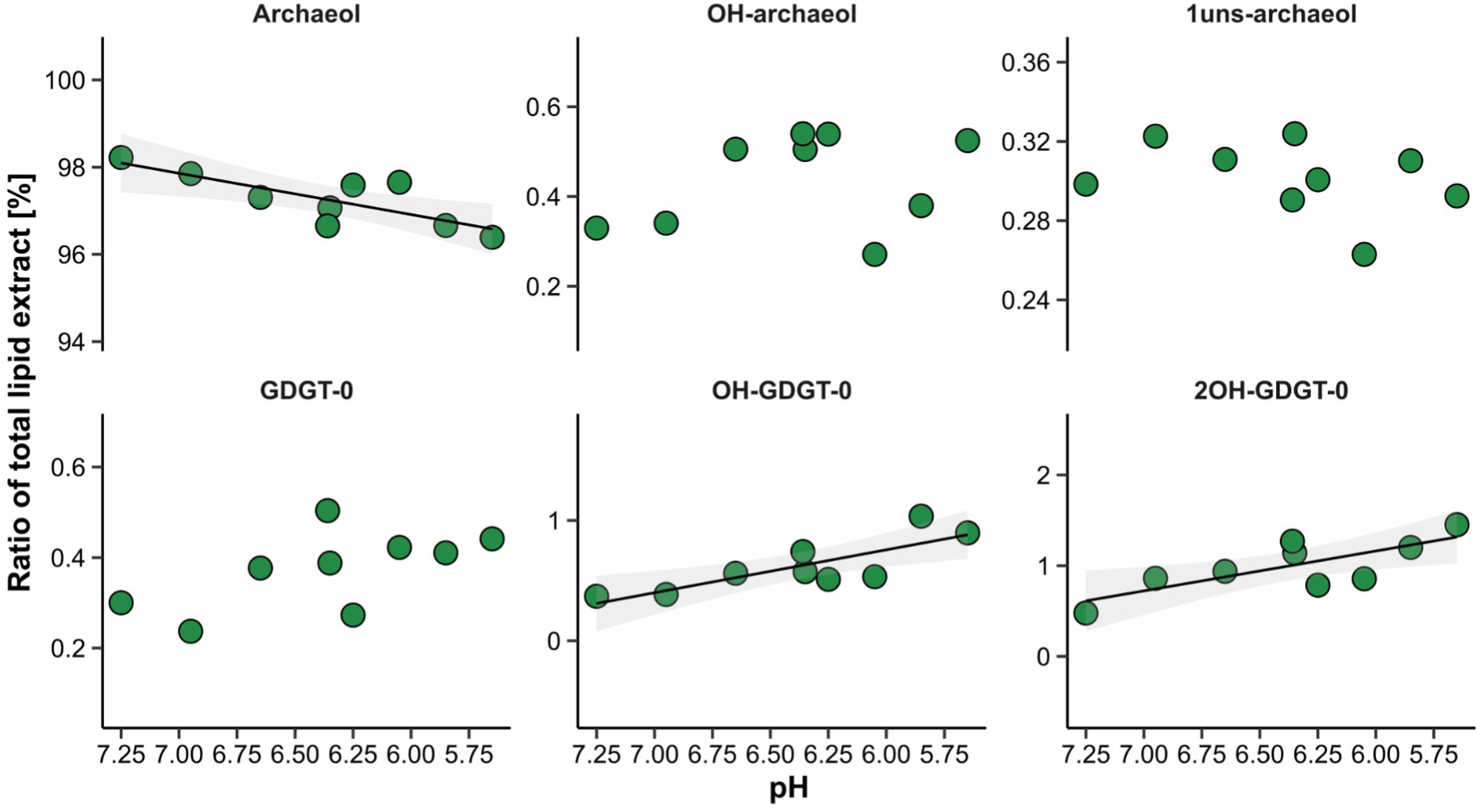
The relative fraction of core lipids extracted from the biomass of ‘*Ca*. M. vercellensis’. Biomass was sampled at the end of each bioreactor batch incubation. The control sample represents biomass before the start of acidification experiment. Solid lines indicate significant relationships (p < 0.05) with 0.95 confidence interval in grey.

In contrast to many studies on monolayer-forming lipids, how archaea with predominantly bilayer-forming lipids adapt to low pH has not been studied previously. Analysis of intact polar lipids (IPLs) of ‘*Ca*. M. vercellensis’ showed a range of archaeols and hydroxyarchaeols with different headgroups, which were grouped based on the charge of their headgroups: neutral, anionic, and zwitterionic. Examples of identified IPLs are denoted in Figure 5D, E, F, G, with MS fragmentation spectra of newly described IPLs shown in Supplementary Figure 5, and their full list is given in Supplementary Table 1. The summed relative abundance of IPLs at the level of these three groups showed that the majority of IPLs are zwitterionic or anionic lipids (44.88 ± 2.81% and 31.13 ± 1.84%, respectively). The change in the fraction of total IPLs for these two groups showed a significant trend with acidification of the bioreactor medium (Figure 7). While anionic IPLs (phosphatidylinositol archaeol and phosphatidylinositol hydroxyarchaeol) were positively correlated with pH (β = 8.95, p < 0.01), zwitterionic IPLs showed strong negative correlation with pH (β = -15.60, p < 0.001). Two IPLs were responsible for the observed increase of zwitterionic IPLs with acidification: phosphatidylethanolamine-1uns-hydroxyarchaeol and phosphatidylethanolamine-archaeol (β = -12.13 and -1.77, respectively, p < 0.001, Supplementary Figure 4, Supplementary Table 2). The fraction of neutral IPLs (mainly diglycosyl archaeol and diglycosyl hydroxyarchaeol) showed a slight increase with pH (β = 2.75, p < 0.01, 10.29 ± 0.56%).

**Figure 7.**
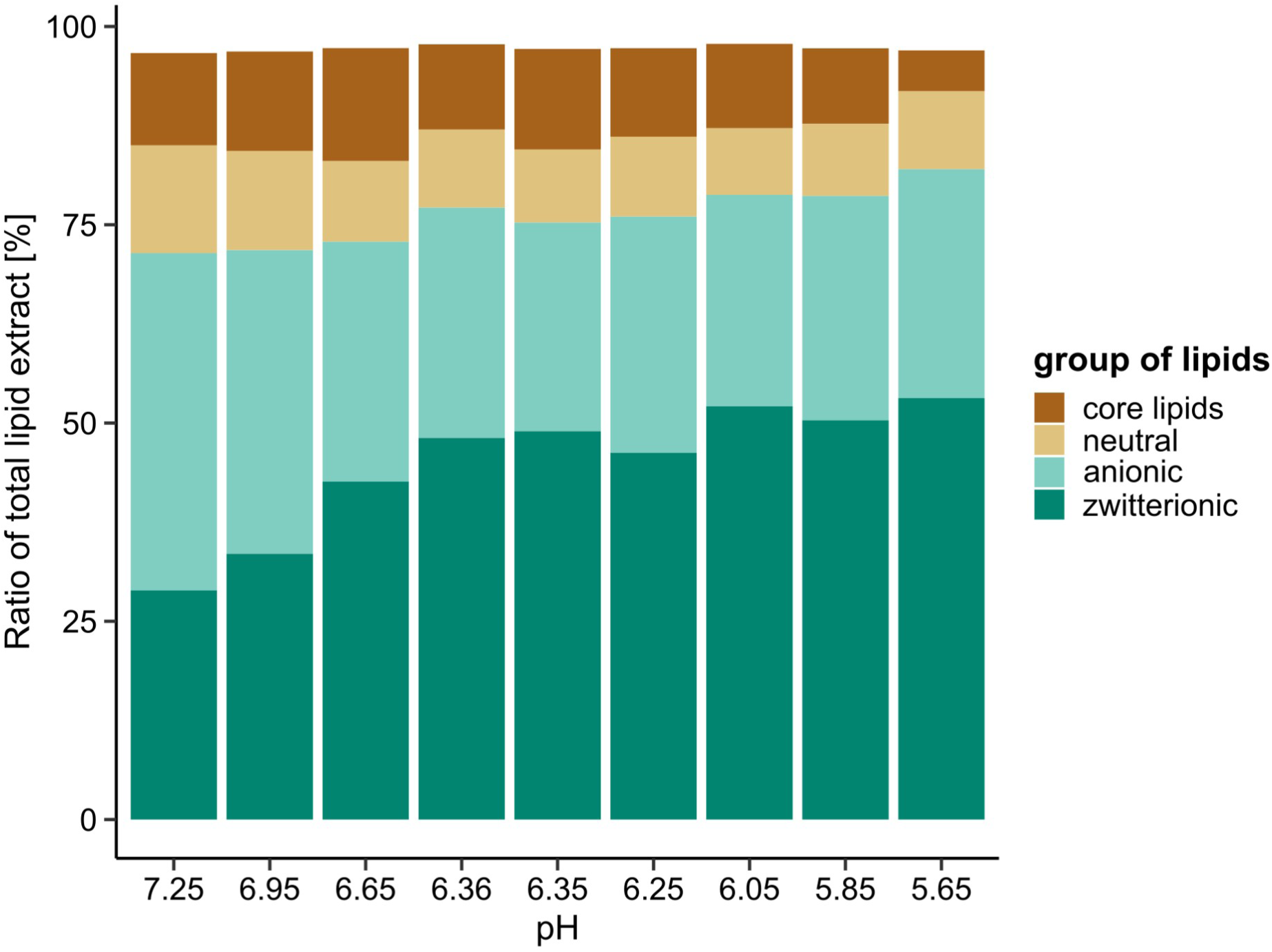
The relative fraction of intact polar lipids extracted from the biomass of ‘*Ca*. M. vercellensis’. Detected lipids are grouped based on their properties given in Supplementary Table 1. The biomass samples were obtained at the end of the respective bioreactor batch incubations at the pH displayed on the X axis.

Based on these results, we suggest that membrane lipid remodelling through an increased proportion of zwitterionic IPLs represents an adaptation mechanism by ‘*Ca*. M. vercellensis’ in response to low pH. IPL adaptation of archaea to changes in pH has only been studied in a few species, such as *T. acidophilum* (Shimada et al., 2008) and *N. maritimus* (Elling et al., 2015). In both studies higher relative abundances of multiple sugar-containing IPLs were observed at lower pH values, while the charge state of lipids has not been investigated and discussed. However, analogies can be drawn to the similar IPL headgroups found in bacteria and their adaptation mechanisms. In bacteria, IPL headgroup composition, and thereby membrane lipid charge, is modulated to retain membrane fluidity, proton permeability, and hydration among other factors (Zhang and Rock, 2008). A decrease in extracellular pH, i.e., an increase in extracellular H^+^/H_3_O^+^ concentration, may impact surface properties of lipid membranes and reduce the efficiency of proton translocation across the membrane. Although both zwitterionic and anionic IPLs are needed for proper membrane functioning in bacteria (Zhang and Rock, 2008), an increase in zwitterionic IPLs may be beneficial under low pH conditions. The small headgroup size and conical shape of phosphatidylethanolamine (PE) IPLs leads to a tight packing of the membrane (Murzyn et al., 2005), which may decrease H^+^ permeability. By contrast, the anionic, phosphatidylinositol (PI) IPLs of ‘*Ca*. M. vercellensis’ are bulky and thus result in a looser packing of the membrane (Peng et al., 2012). Although both PE- and PI-containing IPLs can form hydrogen bonds with each other (Van Den Brink-van Der Laan et al., 2004; Graber et al., 2018), membranes with increased amounts of PE-containing IPLs may still retain high levels of hydrogen bonding through the PE-PE bonding (Hauser et al., 1981; Boggs, 1987), thereby decreasing H^+^ permeability. Still, some amount of PI-containing IPLs or other IPLs are required as the conical shape of PE induces curvature stress (Van Den Brink-van Der Laan et al., 2004) and membranes solely composed of PE-containing IPLs would not form stable bilayers (Cullis and De Kruijff, 1979). Finally, zwitterionic IPLs are less sensitive to H_3_O^+^ binding and dehydration compared to anionic IPLs, preserving membrane hydration at low pH (Abhinav et al., 2022).

We therefore suggest that ‘*Ca*. M. vercellensis’ adapts to low pH by controlling proton permeability and hydration of its membrane through changes in lipid headgroup composition, similar to adaptive mechanisms known from bacteria. It remains unconstrained how this adaptive mechanism is regulated in ‘*Ca*. M. vercellensis’. In bacteria, biosynthesis of anionic vs. zwitterionic IPLs may be regulated at the phosphatidic acid branching point (Zhang and Rock, 2008). In archaea, the biosynthesis of IPL headgroups is largely unconstrained but it is thought that CDP-archaeol is a central branching point (Morii and Koga, 2003; Morii et al., 2009) and therefore the upregulation of phosphatidylethanolamine archaeol biosynthesis could directly impact the biosynthesis of the phosphatidylinositol-based anionic IPLs of ‘*Ca*. M. vercellensis’.

### Biotechnological and environmental implications

Recent studies have been focusing on the occurrence and application of ‘*Ca*. Methanoperedens’ in engineered systems like wastewater treatment plants or landfills. In this study, we show a wider pH activity range which might allow usage of ‘*Ca*. Methanoperedens’ as bioactive agent in CH_4_-contaminated wastewater streams at lower pH values (Liu et al., 2020) or landfill cover soils (Li et al., 2024). For example, anaerobic oxidation of methane in laboratory scale bioreactors has been shown to achieve high nitrogen removal from synthetic wastewater (Fan et al., 2020). Similar laboratory setups often operate at stable pH (Liu et al., 2020, 2023); the microbial community in real-world applications is, however, subjected to dynamic water influent. Activity at low pH would also allow for survival and CH _4_ emission offsetting in wetland and peatland habitats containing nitrate, nitrite, iron, or manganese (Segarra et al., 2015). The latter habitat additionally contains high contents of humic compounds which have been shown to support AOM as regenerable electron acceptors (Klüpfel et al., 2014; Valenzuela et al., 2017; Bai et al., 2019). AOM is further supported by biochar amendment of anoxic habitats like landfills or paddy soils, where this treatment has been applied as a CH_4_ mitigation strategy (Zhang et al., 2019). However, AOM in acidic environments remains understudied (Gupta et al., 2013), and further research is needed to quantify its contribution to CH_4_ turnover in these habitats.

‘*Ca*. M. vercellensis’ has previously been shown to adapt to osmotic stress through the expression of salt-stress genes leading to osmolyte-mediated adjustment of cytoplasmic osmotic pressure, accompanied by utilization of the storage polymer polyhydroxyalkanoate (Echeveste Medrano et al., 2024). Another adaptation of freshwater ‘*Ca*. Methanoperedens’ is represented by tolerance against long-term sulfide exposure (Echeveste Medrano et al., 2025a). Here, we describe its potential to adapt to low pH via remodelling of membrane lipid composition and granular structure. Together, these mechanisms allow methane oxidation under low pH conditions, thereby expanding the environmental niche within which ‘ *Ca*. Methanoperedens’ can influence CH_4_ fluxes.

## Data Records

The raw data used in this paper are available via Zenodo repository.

## Code Availability

Processing scripts are deposited at a public repository https://github.com/TlaskalV/mper_ph_adaptation.

## Acknowledgements

The authors thank Guylaine Nuijten for helping with bioreactor setup, Robert S. Jansen for valuable discussion about annotation of the data, and Anja Engel for access to UPLC-Orbitrap-MS/MS.

## Funding

This work was supported by the SIAM Gravitation Grant (024.002.002) and a VIDI Talent Grant (VI.Vidi.223.012) by the Dutch Science Foundation to CW. It was furthermore supported by an ERC Consolidator grant (electroANME, grant number 101229409) to CW. VT was supported by the Czech Science Foundation grant number 23-07434O. FJE and XZ were supported by Deutsche Forschungsgemeinschaft grants 441217575 and 527682349. The authors used services of the Czech-BioImaging research infrastructure, specifically the Light Microscopy Core Facility, Biology Centre CAS, supported by project LM2023050 funded by the Ministry of Education, Youth and Sports of the Czech Republic, with the instrumental equipment co-financed by the European Union.

## Competing interests statement

The authors declare no competing financial interest.

## CRediT author statement

VT: conceptualization, data curation, formal analysis, visualisation, writing - original draft, writing - review & editing;

RAE: methodology, formal analysis, writing - review & editing;

WW: methodology, formal analysis, writing - review & editing;

XZ: methodology, formal analysis, writing - review & editing;

MW: methodology, writing - review & editing;

MJEM: methodology, writing - review & editing;

KWB: formal analysis, writing - review & editing;

FE: methodology, formal analysis, writing - review & editing;

CUW: conceptualization, funding acquisition, formal analysis, writing - review & editing.

---

**Supplementary Figure 1.**
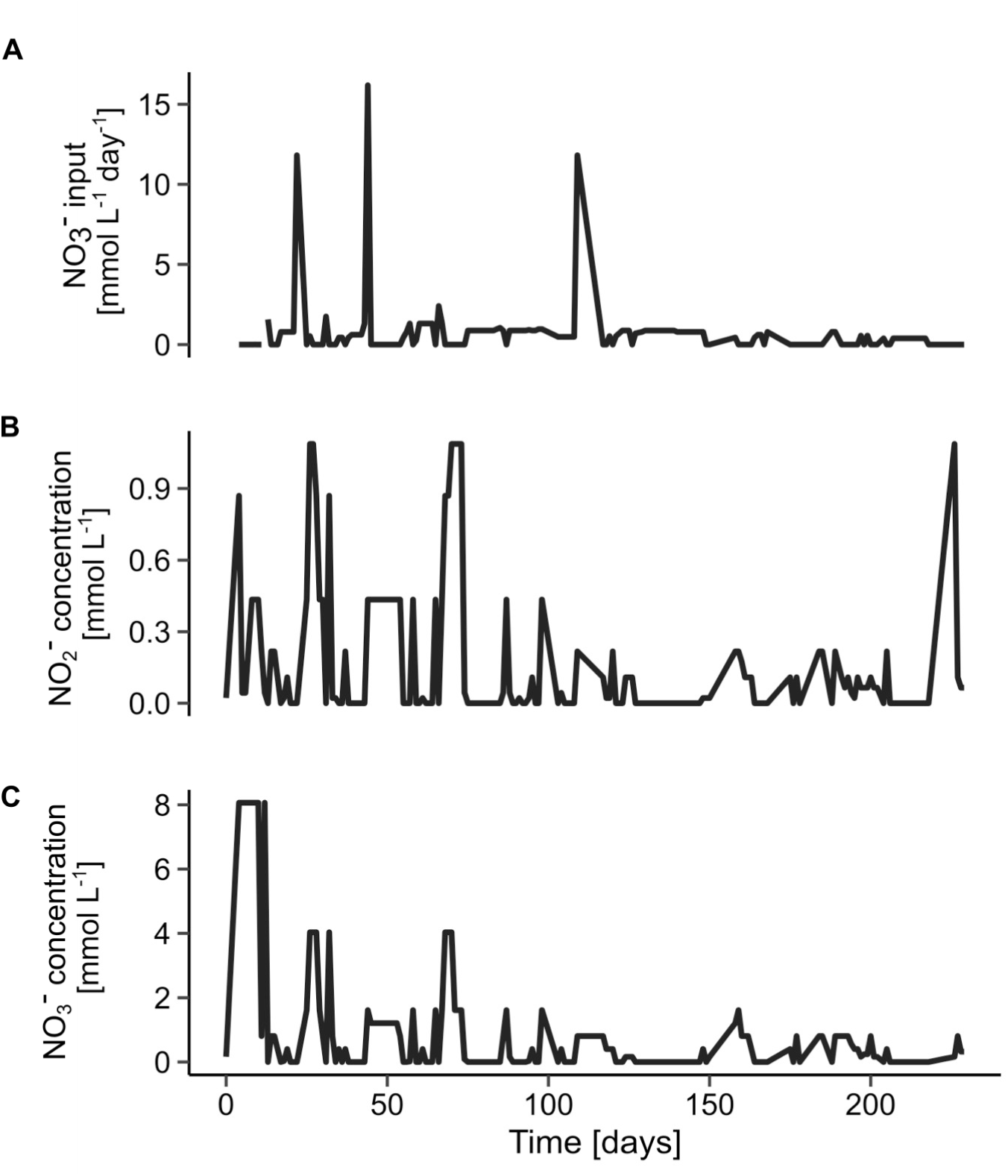
The characteristics of the bioreactor with the enrichment culture of ‘*Ca*. M. vercellensis’ during the 225 days of the acidification experiment. Nitrate added to the bioreactor (in mmol day^-1^, **A**) and concentrations of nitrite (**B**) and nitrate (**C**) measured *in situ* in the bioreactor over the course of the acidification experiment (both in mmol L^-1^).

**Supplementary Figure 2.**
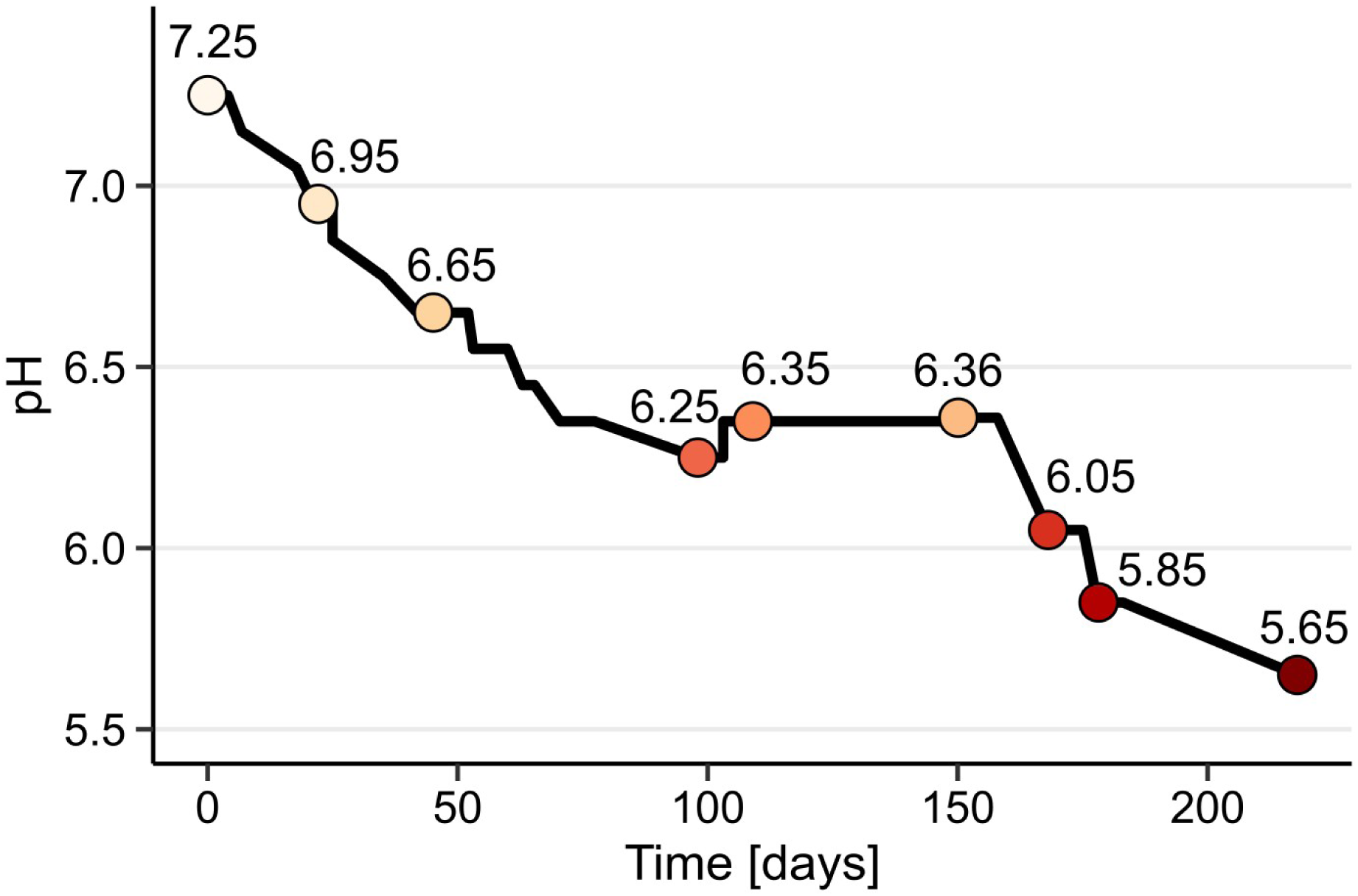
The decrease of pH of bioreactor medium with ‘*Ca*. M. vercellensis’ enrichment culture. Individual points represent timepoints at the beginning of bioreactor batch incubations. The numbers denote pH values.

**Supplementary Figure 3.**
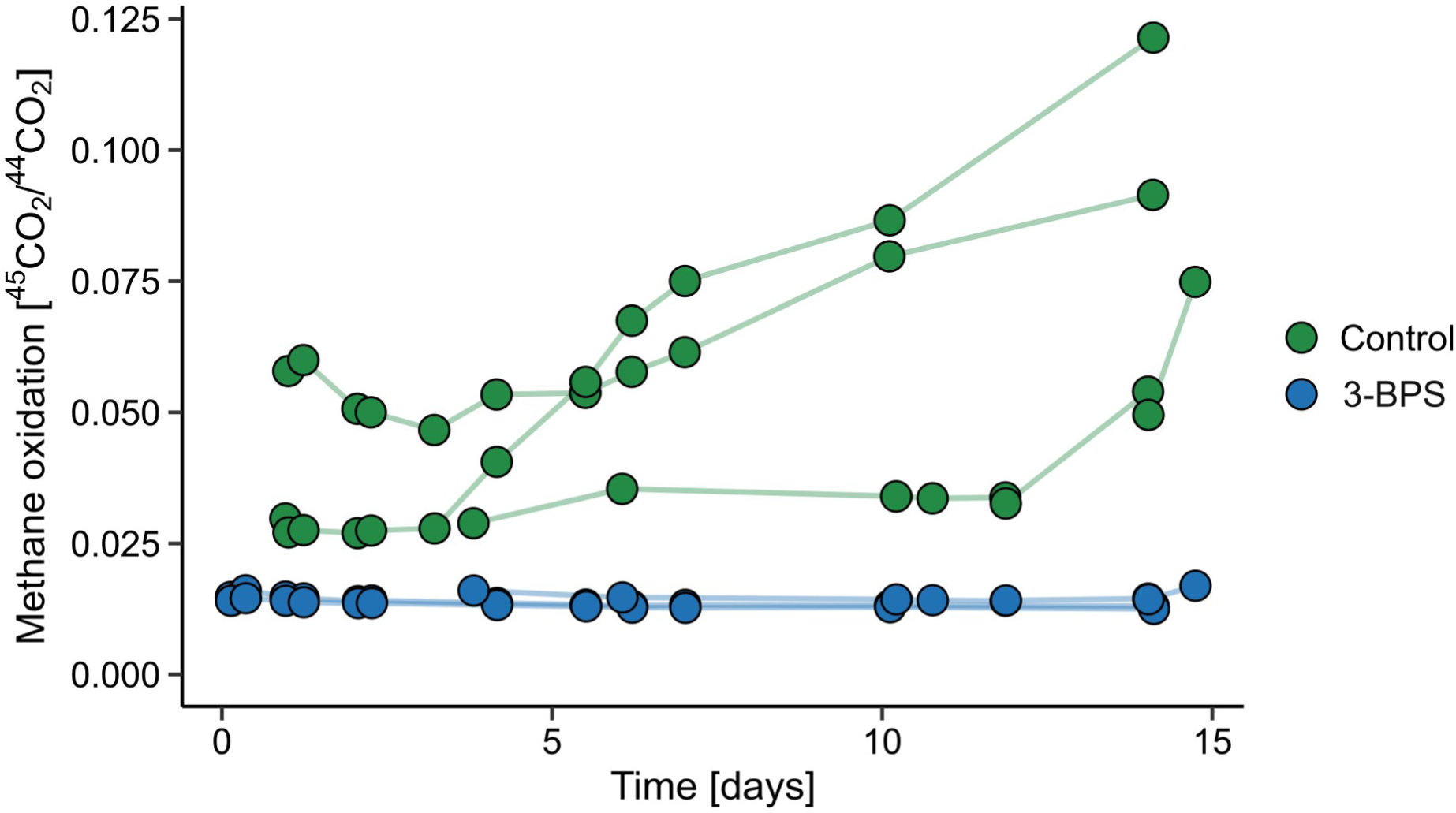
The inhibition of ‘*Ca*. M. vercellensis’ by 3-bromopropanesulfonic acid (3-BPS). ^45^CO_2_/^44^CO_2_ ratio in the headspace of batch incubations with 3-BPS (n = 3 biological replicates) and control bottles (n = 3 biological replicates) was used to assess oxidation of ^13^C-CH₄. Replicates are displayed separately to show the variation of individual batch incubation assays.

**Supplementary Figure 4.**
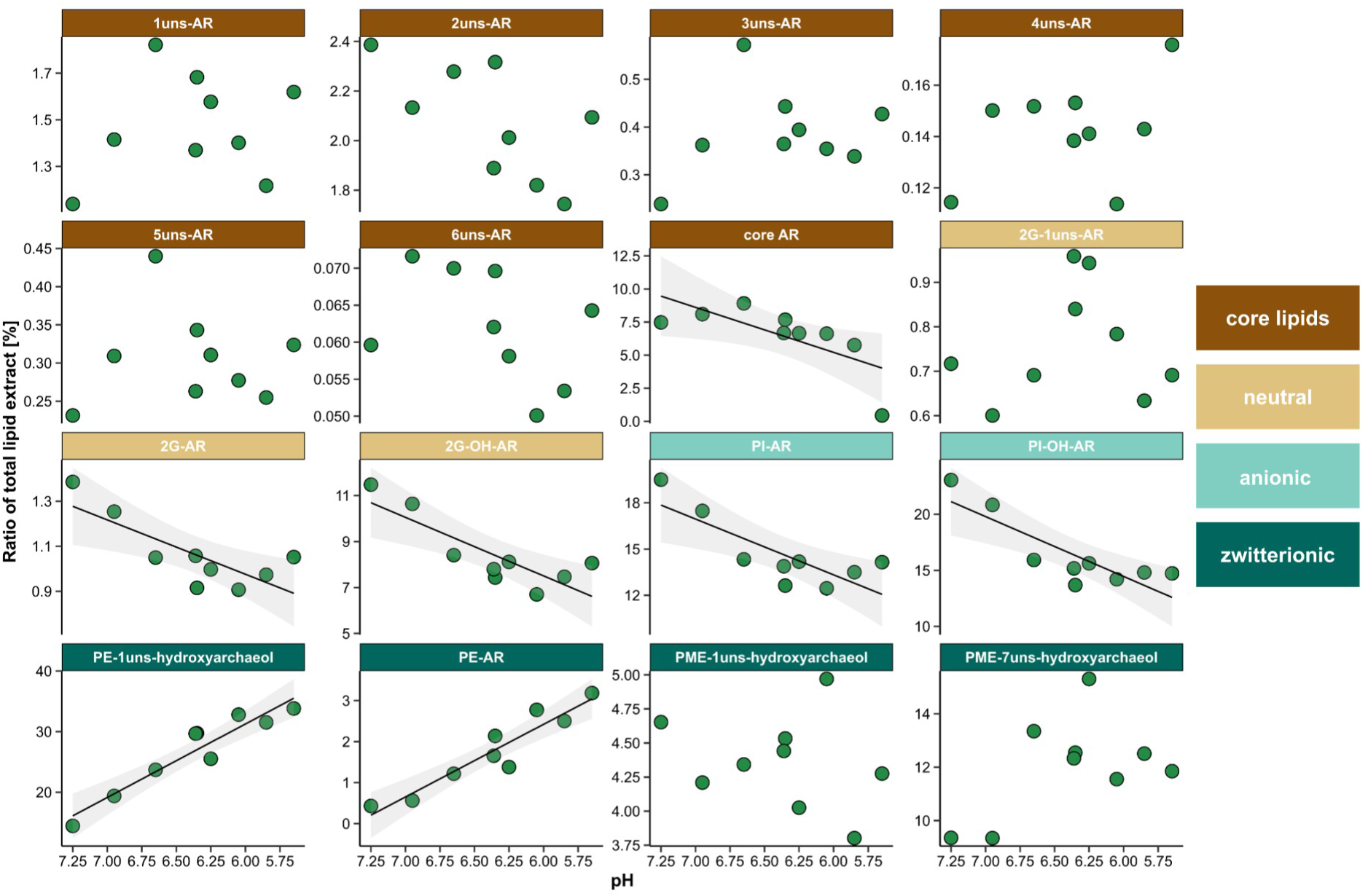
The relative fraction of intact polar lipids extracted from the biomass of ‘*Ca*. M. vercellensis’. Intact polar lipids are grouped and coloured based on the charge of their headgroups: neutral, anionic, and zwitterionic. Core lipids were extracted together with intact polar lipids and were included in the visualisation. The biomass samples were obtained at the end of respective bioreactor batch incubations, solid lines indicate significant relationships (p < 0.05) with 0.95 confidence interval in grey.

**Supplementary Figure 5.**
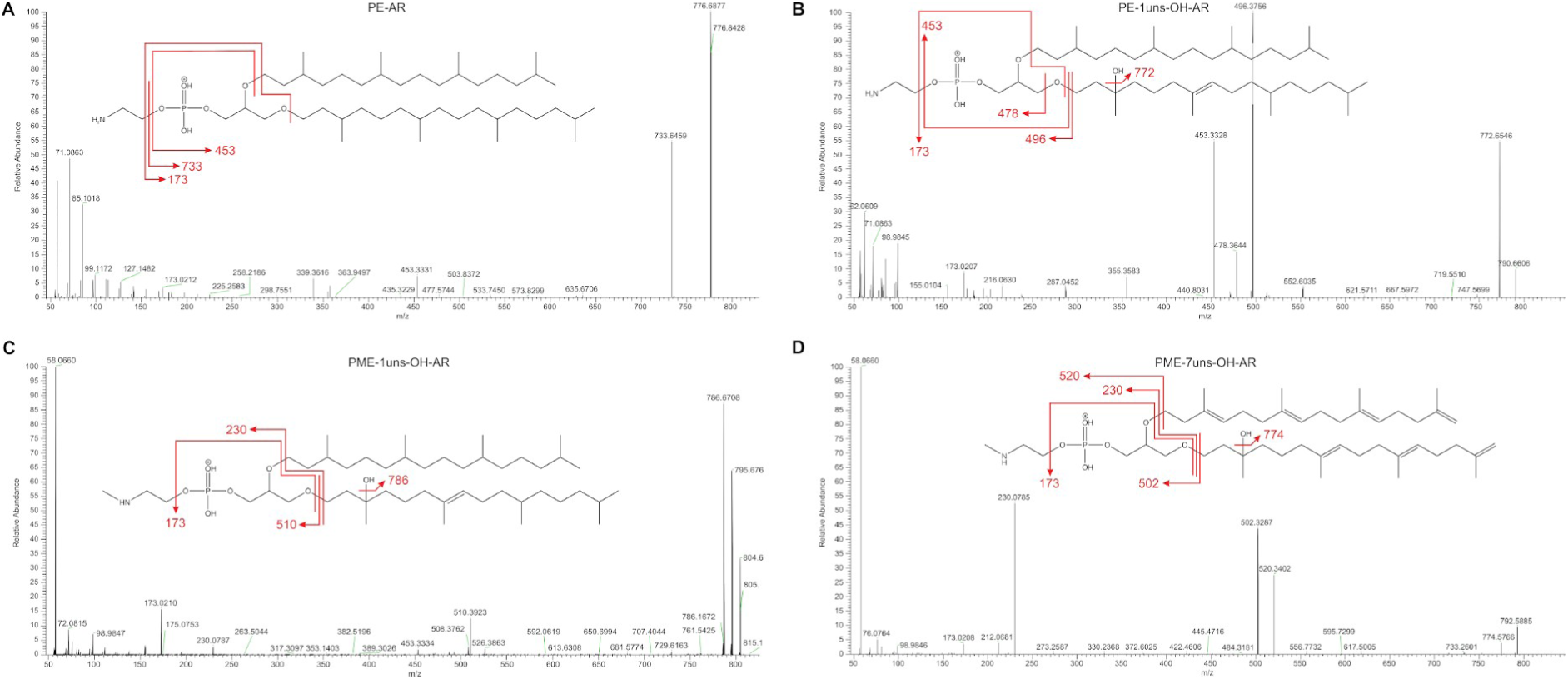
MS^2^ fragmentation spectra of part of the identified archaeols. The spectra of phosphatidylethanolamine-archaeol (**A**), phosphatidylethanolamine-1uns-hydroxyarchaeol (**B**), phosphatidylmethylethanolamine-1uns-hydroxyarchaeol (**C**), and phosphatidylmethylethanolamine-7uns-hydroxyarchaeol (**D**) have not been described before and were detected in intact polar lipid extract from the biomass of ‘*Ca*. M. vercellensis’.

## Notes

### Competing Interest Statement

The authors have declared no competing interest.

## References

1. Abhinav, Jurkiewicz P., Hof, M., Allolio, C., and Sýkora, J. (2022). Modulation of anionic lipid bilayers by specific interplay of protons and calcium ions. Biomolecules 12, 1894. doi: 10.3390/biom12121894

2. Amann, R. I., Binder, B. J., Olson, R. J., Chisholm, S. W., Devereux, R., and Stahl, D. A. (1990). Combination of 16S rRNA-targeted oligonucleotide probes with flow cytometry for analyzing mixed microbial populations. Appl. Environ. Microbiol. 56, 1919–1925. doi: 10.1128/aem.56.6.1919-1925.1990

3. Angel, R., Matthies, D., and Conrad, R. (2011). Activation of methanogenesis in arid biological soil crusts despite the presence of oxygen. PLoS ONE 6, e20453. doi: 10.1371/journal.pone.0020453

4. Angel, R., Petrova, E., and Lara-Rodriguez, A. (2021). Total Nucleic Acids Extraction from Soil v6. doi: 10.17504/protocols.io.bwxcpfiw

5. Arshad, A., Speth, D. R., de Graaf, R. M., Camp, H. J. M. O. D., Jetten, M. S. M., and Welte, C. U. (2015). A metagenomics-based metabolic model of nitrate-dependent anaerobic oxidation of methane by *Methanoperedens*-like Archaea. Front. Microbiol. 6, 1423. doi: 10.3389/fmicb.2015.01423

6. Bai, Y.-N., Wang, X.-N., Wu, J., Lu, Y.-Z., Fu, L., Zhang, F., et al. (2019). Humic substances as electron acceptors for anaerobic oxidation of methane driven by ANME-2d. Water Res. 164, 114935. doi: 10.1016/j.watres.2019.114935

7. Barney, M., Hopple, A. M., Gregory, L. L., Keller, J. K., and Bridgham, S. D. (2024). Anaerobic oxidation of methane mitigates net methane production and responds to long-term experimental warming in a northern bog. Soil Biol. Biochem. 190, 109316. doi: 10.1016/j.soilbio.2024.109316

8. Berger, S., Frank, J., Dalcin Martins, P., Jetten, M. S. M., and Welte, C. U. (2017). High-quality draft genome sequence of “*Candidatus* Methanoperedens sp.” strain BLZ2, a nitrate-reducing anaerobic methane-oxidizing archaeon enriched in an anoxic bioreactor. Genome Announc. 5, e01159–17. doi: 10.1128/genomeA.01159-17

9. Bligh, E. G., and Dyer, W. J. (1959). A rapid method of total lipid extraction and purification. Can. J. Biochem. Physiol. 37, 911–917. doi: 10.1139/o59-099

10. Blum, L. N., Colman, D. R., Eloe-Fadrosh, E. A., Kellom, M., Boyd, E. S., Zhaxybayeva, O., et al. (2023). Distribution and abundance of tetraether lipid cyclization genes in terrestrial hot springs reflect pH. Environ. Microbiol. 25, 1644–1658. doi: 10.1111/1462-2920.16375

11. Boetius, A., Ravenschlag, K., Schubert, C. J., Rickert, D., Widdel, F., Gleseke, A., et al. (2000). A marine microbial consortium apparently mediating anaerobic oxidation methane. Nature 407, 623–626. doi: 10.1038/35036572

12. Boggs, J. M. (1987). Lipid intermolecular hydrogen bonding: influence on structural organization and membrane function. Biochim. Biophys. Acta BBA - Rev. Biomembr. 906, 353–404. doi: 10.1016/0304-4157(87)90017-7

13. Boyd, E. S., Hamilton, T. L., Wang, J., He, L., and Zhang, C. L. (2013). The role of tetraether lipid composition in the adaptation of thermophilic archaea to acidity. Front. Microbiol. 4. doi: 10.3389/fmicb.2013.00062

14. Boyd, E. S., Pearson, A., Pi, Y., Li, W.-J., Zhang, Y. G., He, L., et al. (2011). Temperature and pH controls on glycerol dibiphytanyl glycerol tetraether lipid composition in the hyperthermophilic crenarchaeon *Acidilobus sulfurireducens*. Extremophiles 15, 59–65. doi: 10.1007/s00792-010-0339-y

15. Cai, C., Leu, A. O., Xie, G.-J., Guo, J., Feng, Y., Zhao, J.-X., et al. (2018). A methanotrophic archaeon couples anaerobic oxidation of methane to Fe(III) reduction. ISME J. 12, 1929–1939. doi: 10.1038/s41396-018-0109-x

16. Cai, C., Shi, Y., Guo, J., Tyson, G. W., Hu, S., and Yuan, Z. (2019). Acetate Production from Anaerobic Oxidation of Methane via Intracellular Storage Compounds. Environ. Sci. Technol. 53, 7371–7379. doi: 10.1021/acs.est.9b00077

17. Chong, P. L.-G. (2010). Archaebacterial bipolar tetraether lipids: Physico-chemical and membrane properties. Chem. Phys. Lipids 163, 253–265. doi: 10.1016/j.chemphyslip.2009.12.006

18. Cobban, A., Zhang, Y., Zhou, A., Weber, Y., Elling, F. J., Pearson, A., et al. (2020). Multiple environmental parameters impact lipid cyclization in *Sulfolobus acidocaldarius*. Environ. Microbiol. 22, 4046–4056. doi: 10.1111/1462-2920.15194

19. Cullis, P. R., and De Kruijff, B. (1979). Lipid polymorphism and the functional roles of lipids in biological membranes. Biochim. Biophys. Acta BBA - Rev. Biomembr. 559, 399–420. doi: 10.1016/0304-4157(79)90012-1

20. Echeveste Medrano, M. J., Lee, S., De Graaf, R., Holohan, B. C., Sánchez-Andrea, I., Jetten, M. S. M., et al. (2025a). Physiological stress response to sulfide exposure of freshwater anaerobic methanotrophic archaea. Environ. Sci. Technol., acs.est.4c12489. doi: 10.1021/acs.est.4c12489

21. Echeveste Medrano, M. J., Leu, A. O., Pabst, M., Lin, Y., Mcllroy, S. J., Tyson, G. W., et al. (2024). Osmoregulation in freshwater anaerobic methane oxidizing archaea under salt stress. *ISME J.*, wrae137. doi: 10.1093/ismejo/wrae137

22. Echeveste Medrano, M. J., Su, G., Blattner, L. A., Leão, P., Sánchez-Andrea, I., Jetten, M. S. M., et al. (2025b). Methanotrophic flexibility of “*Ca*. Methanoperedens” and its interactions with sulphate-reducing bacteria in the sediment of meromictic Lake Cadagno. Environ. Microbiol. 27. doi: 10.1111/1462-2920.70133

23. Egas, R. A., Lin, H., Leu, A. O., Tyson, G. W., McIlroy, S. J., and Welte, C. U. (2025). Carbon monoxide metabolism in freshwater anaerobic methanotrophic archaea. doi: 10.1101/2025.09.16.676500

24. Elbers, B. (2020). tidylog: Logging for “dplyr” and “tidyr” Functions. Available at: https://CRAN.R-project.org/package=tidylog

25. Elling, F. J., Könneke, M., Lipp, J. S., Becker, K. W., Gagen, E. J., and Hinrichs, K.-U. (2014). Effects of growth phase on the membrane lipid composition of the thaumarchaeon *Nitrosopumilus maritimus* and their implications for archaeal lipid distributions in the marine environment. Geochim. Cosmochim. Acta 141, 579–597. doi: 10.1016/j.gca.2014.07.005

26. Elling, F. J., Könneke, M., Mußmann, M., Greve, A., and Hinrichs, K.-U. (2015). Influence of temperature, pH, and salinity on membrane lipid composition and TEX_86_ of marine planktonic thaumarchaeal isolates. Geochim. Cosmochim. Acta 171, 238–255. doi: 10.1016/j.gca.2015.09.004

27. Ettwig, K. F., Zhu, B., Speth, D., Keltjens, J. T., Jetten, M. S. M., and Kartal, B. (2016). Archaea catalyze iron-dependent anaerobic oxidation of methane. Proc. Natl. Acad. Sci. 113, 12792–12796. doi: 10.1073/pnas.1609534113

28. Fan, S.-Q., Xie, G.-J., Lu, Y., Liu, B.-F., Xing, D.-F., Han, H.-J., et al. (2020). Granular sludge coupling nitrate/nitrite dependent anaerobic methane oxidation with anammox: From proof-of-concept to high rate nitrogen removal. Environ. Sci. Technol. 54, 297– 305. doi: 10.1021/acs.est.9b02528

29. Feyhl-Buska, J., Chen, Y., Jia, C., Wang, J.-X., Zhang, C. L., and Boyd, E. S. (2016). Influence of growth phase, pH, and temperature on the abundance and composition of tetraether lipids in the thermoacidophile *Picrophilus torridus*. Front. Microbiol. 7. doi: 10.3389/fmicb.2016.01323

30. Frank, J., Zhang, X., Marcellin, E., Yuan, Z., and Hu, S. (2023). Salinity effect on an anaerobic methane- and ammonium-oxidising consortium: Shifts in activity, morphology, osmoregulation and syntrophic relationship. Water Res. 242, 120090. doi: 10.1016/j.watres.2023.120090

31. Graber, Z. T., Thomas, J., Johnson, E., Gericke, A., and Kooijman, E. E. (2018). Effect of H-Bond Donor Lipids on Phosphatidylinositol-3,4,5-Trisphosphate Ionization and Clustering. Biophys. J. 114, 126–136. doi: 10.1016/j.bpj.2017.10.029

32. Gupta, V., Smemo, K. A., Yavitt, J. B., Fowle, D., Branfireun, B., and Basiliko, N. (2013). Stable isotopes reveal widespread anaerobic methane oxidation across latitude and peatland type. Environ. Sci. Technol. 47, 8273–8279. doi: 10.1021/es400484t

33. Haroon, M. F., Hu, S., Shi, Y., Imelfort, M., Keller, J., Hugenholtz, P., et al. (2013). Anaerobic oxidation of methane coupled to nitrate reduction in a novel archaeal lineage. Nature 500, 567–570. doi: 10.1038/nature12375

34. Harpenslager, S. F., Van Dijk, G., Boonman, J., Weideveld, S. T. J., Van De Riet, B. P., Hefting, M. M., et al. (2024). Rewetting drained peatlands through subsoil infiltration stabilises redox-dependent soil carbon and nutrient dynamics. Geoderma 442, 116787. doi: 10.1016/j.geoderma.2024.116787

35. Hauser, H., Pascher, I., Pearson, R. H., and Sundell, S. (1981). Preferred conformation and molecular packing of phosphatidylethanolamine and phosphatidylcholine. Biochim. Biophys. Acta BBA - Rev. Biomembr. 650, 21–51. doi: 10.1016/0304-4157(81)90007-1

36. Hopmans, E. C., Schouten, S., and Sinninghe Damsté, J. S. (2016). The effect of improved chromatography on GDGT-based palaeoproxies. Org. Geochem. 93, 1–6. doi: 10.1016/j.orggeochem.2015.12.006

37. IPCC (2023). Climate Change 2021 – The Physical Science Basis: Working Group I Contribution to the Sixth Assessment Report of the Intergovernmental Panel on Climate Change., 1st Edn. Cambridge University Press. doi: 10.1017/9781009157896

38. Kang, H., Kwon, M. J., Kim, S., Lee, S., Jones, T. G., Johncock, A. C., et al. (2018). Biologically driven DOC release from peatlands during recovery from acidification. Nat. Commun. 9, 3807. doi: 10.1038/s41467-018-06259-1

39. Kletzin, A., Heimerl, T., Flechsler, J., Van Niftrik, L., Rachel, R., and Klingl, A. (2015). Cytochromes c in Archaea: distribution, maturation, cell architecture, and the special case of *Ignicoccus hospitalis*. Front. Microbiol. 6. doi: 10.3389/fmicb.2015.00439

40. Klüpfel, L., Piepenbrock, A., Kappler, A., and Sander, M. (2014). Humic substances as fully regenerable electron acceptors in recurrently anoxic environments. Nat. Geosci. 7, 195–200. doi: 10.1038/ngeo2084

41. Krause, S. J. E., and Treude, T. (2021). Deciphering cryptic methane cycling: Coupling of methylotrophic methanogenesis and anaerobic oxidation of methane in hypersaline coastal wetland sediment. Geochim. Cosmochim. Acta 302, 160–174. doi: 10.1016/j.gca.2021.03.021

42. Kurth, J. M., Smit, N. T., Berger, S., Schouten, S., Jetten, M. S. M., and Welte, C. U. (2019). Anaerobic methanotrophic archaea of the ANME-2d clade feature lipid composition that differs from other ANME archaea. FEMS Microbiol. Ecol. 95, fiz082. doi: 10.1093/femsec/fiz082

43. Lan, X., Thoning, K., Dlugokencky, E., and NOAA Global Monitoring Laboratory (2025). Trends in globally-averaged CH_4_, N_2_O, and SF6 determined from NOAA Global Monitoring Laboratory measurements. doi: 10.15138/P8XG-AA10

44. Leu, A. O., Cai, C., McIlroy, S. J., Southam, G., Orphan, V. J., Yuan, Z., et al. (2020). Anaerobic methane oxidation coupled to manganese reduction by members of the *Methanoperedenaceae*. ISME J. 14, 1030–1041. doi: 10.1038/s41396-020-0590-x

45. Li, R., Xi, B., Wang, X., Li, Y., Yuan, Y., and Tan, W. (2024). Anaerobic oxidation of methane in landfill and adjacent groundwater environments: Occurrence, mechanisms, and potential applications. Water Res. 255, 121498. doi: 10.1016/j.watres.2024.121498

46. Liu, T., Hu, S., Yuan, Z., and Guo, J. (2023). Simultaneous dissolved methane and nitrogen removal from low-strength wastewater using anaerobic granule-based sequencing batch reactor. Water Res. 242, 120194. doi: 10.1016/j.watres.2023.120194

47. Liu, T., Lim, Z. K., Chen, H., Wang, Z., Hu, S., Yuan, Z., et al. (2020). Biogas-driven complete nitrogen removal from wastewater generated in side-stream partial nitritation. Sci. Total Environ. 745, 141153. doi: 10.1016/j.scitotenv.2020.141153

48. Liu, T., Zhang, X., Hu, S., Yuan, Z., and Guo, J. (2026). Methanoperedenaceae archaea: a 20-year research journey. Nat. Commun. doi: 10.1038/s41467-026-69699-0

49. Makowski, D., Lüdecke, D., Patil, I., Thériault, R., Ben-Shachar, M. S., and Wiernik, B. M. (2023). Automated results reporting as a practical tool to improve reproducibility and methodological best practices adoption. CRAN. Available at: https://easystats.github.io/report/

50. Malcolm, I. A., Gibbins, C. N., Fryer, R. J., Keay, J., Tetzlaff, D., and Soulsby, C. (2014). The influence of forestry on acidification and recovery: Insights from long-term hydrochemical and invertebrate data. Ecol. Indic. 37, 317–329. doi: 10.1016/j.ecolind.2011.12.011

51. McIlroy, S. J., Leu, A. O., Zhang, X., Newell, R., Woodcroft, B. J., Yuan, Z., et al. (2023). Anaerobic methanotroph ‘*Candidatus* Methanoperedens nitroreducens’ has a pleomorphic life cycle. Nat. Microbiol. doi: 10.1038/s41564-022-01292-9

52. Miller, K. E., Lai, C. T., Dahlgren, R. A., and Lipson, D. A. (2019). Anaerobic methane oxidation in high-arctic Alaskan peatlands as a significant control on net CH_4_ fluxes. Soil Syst. 3, 7. doi: 10.3390/soilsystems3010007

53. Morii, H., Kiyonari, S., Ishino, Y., and Koga, Y. (2009). A novel biosynthetic pathway of archaetidyl-*myo*-inositol via archaetidyl-*myo*-inositol phosphate from CDP-archaeol and D-glucose 6-phosphate in methanoarchaeon *Methanothermobacter thermautotrophicus* cells. J. Biol. Chem. 284, 30766–30774. doi: 10.1074/jbc.M109.034652

54. Morii, H., and Koga, Y. (2003). CDP-2,3-di-O-geranylgeranyl-*sn*-glycerol:*l*-serine *O*-archaetidyltransferase (archaetidylserine synthase) in the methanogenic archaeon *Methanothermobacter thermautotrophicus*. J. Bacteriol. 185, 1181–1189. doi: 10.1128/JB.185.4.1181-1189.2003

55. Murzyn, K., Róg, T., and Pasenkiewicz-Gierula, M. (2005). Phosphatidylethanolamine-Phosphatidylglycerol Bilayer as a Model of the Inner Bacterial Membrane. Biophys. J. 88, 1091–1103. doi: 10.1529/biophysj.104.048835

56. Nweze, J. A., Tláskal, V., Wutkowska, M., Meador, T. B., Picek, T., Urbanová, Z., et al. (2024). Regulators of aerobic and anaerobic methane oxidation in two pristine temperate peatland types. FEMS Microbiol. Ecol. 100, fiae153. doi: 10.1093/femsec/fiae153

57. Ouboter, H. T., Berben, T., Berger, S., Jetten, M. S. M., Sleutels, T., Ter Heijne, A., et al. (2022). Methane-dependent extracellular electron transfer at the bioanode by the anaerobic archaeal methanotroph “*Candidatus* Methanoperedens.” Front. Microbiol. 13, 820989. doi: 10.3389/fmicb.2022.820989

58. Ouboter, H. T., Mesman, R., Sleutels, T., Postma, J., Wissink, M., Jetten, M. S. M., et al. (2024). Mechanisms of extracellular electron transfer in anaerobic methanotrophic archaea. Nat. Commun. 15, 1477. doi: 10.1038/s41467-024-45758-2

59. Peng, A., Pisal, D. S., Doty, A., and Balu-Iyer, S. V. (2012). Phosphatidylinositol induces fluid phase formation and packing defects in phosphatidylcholine model membranes. Chem. Phys. Lipids 165, 15–22. doi: 10.1016/j.chemphyslip.2011.10.002

60. R Core Team (2024). R: A language and environment for statistical computing. Available at: http://www.r-project.org/

61. Raghoebarsing, A. A., Pol, A., Van De Pas-Schoonen, K. T., Smolders, A. J. P., Ettwig, K. F., Rijpstra, W. I. C., et al. (2006). A microbial consortium couples anaerobic methane oxidation to denitrification. Nature 440, 918–921. doi: 10.1038/nature04617

62. Robinson, D., and Hayes, A. (2023). broom: Convert Statistical Objects into Tidy Tibbles. Available at: https://CRAN.R-project.org/package=broom

63. Roy, B., Thiede, R., Simon, S., Kumar, A., Moharana, S., Dey, S., et al. (2025). Elevation controls bacterial branched GDGT-based temperature proxies: A regional to global perspective. Glob. Planet. Change 255, 105101. doi: 10.1016/j.gloplacha.2025.105101

64. Saunois, M., Martinez, A., Poulter, B., Zhang, Z., Raymond, P. A., Regnier, P., et al. (2025). Global methane budget 2000–2020. Earth Syst. Sci. Data 17, 1873–1958. doi: 10.5194/essd-17-1873-2025

65. Seemann, T. (2014). Prokka: Rapid prokaryotic genome annotation. Bioinformatics 30, 2068–2069. doi: 10.1093/bioinformatics/btu153

66. Segarra, K. E. A., Schubotz, F., Samarkin, V., Yoshinaga, M. Y., Hinrichs, K. U., and Joye, S. B. (2015). High rates of anaerobic methane oxidation in freshwater wetlands reduce potential atmospheric methane emissions. Nat. Commun. 6, 7477. doi: 10.1038/ncomms8477

67. Shimada, H., Nemoto, N., Shida, Y., Oshima, T., and Yamagishi, A. (2008). Effects of pH and Temperature on the Composition of Polar Lipids in *Thermoplasma acidophilum* HO-62. J. Bacteriol. 190, 5404–5411. doi: 10.1128/JB.00415-08

68. Smemo, K. A., and Yavitt, J. B. (2011). Anaerobic oxidation of methane: An underappreciated aspect of methane cycling in peatland ecosystems? Biogeosciences 8, 779–793. doi: 10.5194/bg-8-779-2011

69. Stahl, D. A. (1991). “Development and application of nucleic acid probes in bacterial systematics,” in Sequencing and hybridization techniques in bacterial systematics, (Chichester, England: John Wiley & Sons Ltd.).

70. Steinberg, L. M., and Regan, J. M. (2008). Phylogenetic comparison of the methanogenic communities from an acidic, oligotrophic fen and an anaerobic digester treating municipal wastewater sludge. Appl. Environ. Microbiol. 74, 6663–6671. doi: 10.1128/AEM.00553-08

71. Sturt, H. F., Summons, R. E., Smith, K., Elvert, M., and Hinrichs, K. (2004). Intact polar membrane lipids in prokaryotes and sediments deciphered by high-performance liquid chromatography/electrospray ionization multistage mass spectrometry—new biomarkers for biogeochemistry and microbial ecology. Rapid Commun. Mass Spectrom. 18, 617–628. doi: 10.1002/rcm.1378

72. Tian, W., Petrová, E., Sakai, S., Nweze, J. E., Daebeler, A., and Angel, R. (2026). Cultivation and genomic characterization of novel methanogens from arid desert biocrust. ISME Commun., ycag013. doi: 10.1093/ismeco/ycag013

73. Timmers, P. H. A., Suarez-Zuluaga, D. A., van Rossem, M., Diender, M., Stams, A. J. M., and Plugge, C. M. (2016). Anaerobic oxidation of methane associated with sulfate reduction in a natural freshwater gas source. ISME J. 10, 1400–1412. doi: 10.1038/ismej.2015.213

74. Timmers, P. H. A., Welte, C. U., Koehorst, J. J., Plugge, C. M., Jetten, M. S. M., and Stams, A. J. M. (2017). Reverse methanogenesis and respiration in methanotrophic Archaea. Archaea 2017, 1654237. doi: 10.1155/2017/1654237

75. Urbanová, Z., and Bárta, J. (2016). Effects of long-term drainage on microbial community composition vary between peatland types. Soil Biol. Biochem. 92, 16–26. doi: 10.1016/j.soilbio.2015.09.017

76. Vaksmaa, A., Guerrero-Cruz, S., Van Alen, T. A., Cremers, G., Ettwig, K. F., Lüke, C., et al. (2017). Enrichment of anaerobic nitrate-dependent methanotrophic ‘*Candidatus* Methanoperedens nitroreducens’ archaea from an Italian paddy field soil. Appl. Microbiol. Biotechnol. 101, 7075–7084. doi: 10.1007/s00253-017-8416-0

77. Valenzuela, E. I., and Cervantes, F. J. (2021). The role of humic substances in mitigating greenhouse gases emissions: Current knowledge and research gaps. Sci. Total Environ. 750, 141677. doi: 10.1016/j.scitotenv.2020.141677

78. Valenzuela, E. I., Prieto-Davó, A., López-Lozano, N. E., Hernández-Eligio, A., Vega-Alvarado, L., Juárez, K., et al. (2017). Anaerobic methane oxidation driven by microbial reduction of natural organic matter in a tropical wetland. Appl. Environ. Microbiol. 83, e00645–17. doi: 10.1128/AEM.00645-17

79. Van Den Brink-van Der Laan, E., Antoinette Killian, J., and De Kruijff, B. (2004). Nonbilayer lipids affect peripheral and integral membrane proteins via changes in the lateral pressure profile. Biochim. Biophys. Acta BBA - Biomembr. 1666, 275–288. doi: 10.1016/j.bbamem.2004.06.010

80. Wagner, M., Horn, M., and Daims, H. (2003). Fluorescence in situ hybridisation for the identification and characterisation of prokaryotes. Curr. Opin. Microbiol. 6, 302–309. doi: 10.1016/S1369-5274(03)00054-7

81. Wallenius, A. J., Venetz, J., Zygadlowska, O. M., Lenstra, W. K., Van Helmond, N. A. G. M., Martins, P. D., et al. (2025). A ubiquitous and diverse methanogenic community drives microbial methane cycling in eutrophic coastal sediments. FEMS Microbiol. Ecol. doi: 10.1093/femsec/fiaf075

82. Wallner, G., Amann, R., and Beisker, W. (1993). Optimizing fluorescent in situ hybridization with rRNA-targeted oligonucleotide probes for flow cytometric identification of microorganisms. Cytometry 14, 136–143. doi: 10.1002/cyto.990140205

83. Welte, C. U., Rasigraf, O., Vaksmaa, A., Versantvoort, W., Arshad, A., Op den Camp, H. J. M., et al. (2016). Nitrate- and nitrite-dependent anaerobic oxidation of methane. Environ. Microbiol. Rep. 8, 941–955. doi: 10.1111/1758-2229.12487

84. Wickham, H., Averick, M., Bryan, J., Chang, W., Mcgowan, L. D. A., François, R., et al. (2019). Welcome to the Tidyverse. J. Open Source Softw. 4, 1686. doi: 10.21105/joss.01686

85. Wickham, H., and Bryan, J. (2023). readxl: Read excel files.

86. Wickham, H., Hester, J., and Bryan, J. (2024). readr: Read rectangular text data. Available at: https://CRAN.R-project.org/package=readr

87. Wissink, M., Engilberge, S., Leão, P., Jansen, R. S., Jetten, M. S. M., Royant, A., et al. (2026). Mini-bacterioferritins: Structural insight into a new type of ferritin-like protein from an anaerobic methane-oxidising archaeon. *Commun*. Biol. doi: 10.1038/s42003-026-09796-4

88. Wissink, M., Glodowska, M., Van Der Kolk, M. R., Jetten, M. S. M., and Welte, C. U. (2024). Probing denitrifying anaerobic methane oxidation via antimicrobial intervention: Implications for innovative wastewater management. Environ. Sci. Technol. 58, 6250–6257. doi: 10.1021/acs.est.3c07197

89. Wörmer, L., Lipp, J. S., Schröder, J. M., and Hinrichs, K.-U. (2013). Application of two new LC–ESI–MS methods for improved detection of intact polar lipids (IPLs) in environmental samples. Org. Geochem. 59, 10–21. doi: 10.1016/j.orggeochem.2013.03.004

90. Yoshinaga, M. Y., Kellermann, M. Y., Rossel, P. E., Schubotz, F., Lipp, J. S., and Hinrichs, K. (2011). Systematic fragmentation patterns of archaeal intact polar lipids by high-performance liquid chromatography/electrospray ionization ion-trap mass spectrometry. Rapid Commun. Mass Spectrom. 25, 3563–3574. doi: 10.1002/rcm.5251

91. Yu, Y., Lee, C., Kim, J., and Hwang, S. (2005). Group-specific primer and probe sets to detect methanogenic communities using quantitative real-time polymerase chain reaction. Biotechnol. Bioeng. 89, 670–679. doi: 10.1002/bit.20347

92. Zeng, Z., Liu, X.-L., Farley, K. R., Wei, J. H., Metcalf, W. W., Summons, R. E., et al. (2019). GDGT cyclization proteins identify the dominant archaeal sources of tetraether lipids in the ocean. Proc. Natl. Acad. Sci. 116, 22505–22511. doi: 10.1073/pnas.1909306116

93. Zhang, X., Xia, J., Pu, J., Cai, C., Tyson, G. W., Yuan, Z., et al. (2019). Biochar-mediated anaerobic oxidation of methane. Environ. Sci. Technol. 53, 6660–6668. doi: 10.1021/acs.est.9b01345

94. Zhang, Y.-M., and Rock, C. O. (2008). Membrane lipid homeostasis in bacteria. Nat. Rev. Microbiol. 6, 222–233. doi: 10.1038/nrmicro1839

95. Zhou, A., Weber, Y., Chiu, B. K., Elling, F. J., Cobban, A. B., Pearson, A., et al. (2020). Energy flux controls tetraether lipid cyclization in *Sulfolobus acidocaldarius*. Environ. Microbiol. 22, 343–353. doi: 10.1111/1462-2920.14851

96. Zhu, C., Lipp, J. S., Wörmer, L., Becker, K. W., Schröder, J., and Hinrichs, K.-U. (2013). Comprehensive glycerol ether lipid fingerprints through a novel reversed phase liquid chromatography–mass spectrometry protocol. Org. Geochem. 65, 53–62. doi: 10.1016/j.orggeochem.2013.09.012

